# Suppression of canonical TGF-β signaling enables GATA4 to interact with H3K27me3 demethylase JMJD3 to promote cardiomyogenesis

**DOI:** 10.1101/2020.02.12.945790

**Authors:** Andrew S. Riching, Etienne Danis, Yuanbiao Zhao, Yingqiong Cao, Congwu Chi, Rushita A. Bagchi, Brianna J. Klein, Hongyan Xu, Tatiana G. Kutateladze, Timothy A. McKinsey, Peter M. Buttrick, Kunhua Song

## Abstract

Direct reprogramming of fibroblasts into cardiomyocytes (CMs) represents a promising strategy to regenerate CMs lost after ischemic heart injury. Overexpression of <u>G</u>ATA4, <u>H</u>AND2, <u>M</u>EF2C, <u>T</u>BX5, miR-1, and miR-133 (GHMT2m) along with transforming growth factor beta (TGF-β) inhibition efficiently promotes reprogramming. However, the mechanisms by which TGF-β blockade promotes cardiac reprogramming remain unknown. Here, we identify interactions between the histone H3 lysine 27 trimethylation (H3K27me3) – demethylase JMJD3, the SWI/SNF remodeling complex subunit BRG1, and cardiac transcription factors. Furthermore, canonical TGF-β signaling regulates the interaction between GATA4 and JMJD3. TGF-β activation impairs the ability of GATA4 to bind target genes and prevents demethylation of H3K27 at cardiac gene promoters during cardiac reprogramming. Finally, a mutation in *GATA4* (V267M) exhibits reduced binding to JMJD3 and impaired cardiomyogenesis. Thus, we have identified an epigenetic mechanism wherein canonical TGF-β pathway activation impairs cardiac gene programming by interfering with GATA4-JMJD3 interactions.

## Introduction

Ischemic heart disease (IHD) accounts for nearly 9 million annual deaths worldwide and the mortality rate associated IHD has begun to plateau in developed countries despite advances in clinical intervention (Roth et al., 2017). Ischemic injury causes loss of cardiomyocytes (CMs), which leads to fibrotic remodeling and impaired heart function (Laflamme and Murry, 2005; Mercola et al., 2011; Talman and Ruskoaho, 2016). CMs in the adult heart have limited (<1% annual) turnover (Bergmann et al., 2009), and current clinical therapies, while effective at slowing progression of disease, do not promote regeneration of lost CMs (Hashimoto et al., 2018). The landmark study demonstrating conversion of fibroblasts to induced pluripotent stem cells via transcription factor overexpression (Takahashi et al., 2007; Takahashi and Yamanaka, 2006) inspired the development of transcription factor cocktails that reprogram fibroblasts into other terminally differentiated cell types including neurons (Vierbuchen et al., 2010), hepatocytes (Huang et al., 2011), and CMs (Ieda et al., 2010). Overexpression of cardiac transcription factors GATA4, MEF2C, and TBX5 converted fibroblasts into induced CMs (iCMs), albeit with low efficiency (Ieda et al., 2010). Since this initial report, many studies have focused on optimizing the efficiency of reprogramming by overexpressing additional cardiac transcription factors (Addis et al., 2013; Hirai and Kikyo, 2014; Ifkovits et al., 2014; Song et al., 2012; Zhao et al., 2015), epigenetic modifiers (Christoforou et al., 2013), microRNAs (Muraoka et al., 2014; Nam et al., 2013), or AKT (Zhou et al., 2015). Additionally, knockdown of repressive epigenetic complex components (Dal-Pra et al., 2017; Zhou et al., 2016) or splicing factors (Liu et al., 2017) have also demonstrated improved reprogramming efficiency. Cardiac reprogramming has also been achieved exclusively by microRNA overexpression (Jayawardena et al., 2012) or treatment with small molecules (Cao et al., 2016). More recently, optimized protocols have led to reprogramming efficiencies in excess of 50% (Zhao et al., 2015; Zhou et al., 2015; Zhou et al., 2017; Zhou et al., 2016) and have identified signaling pathways that impair the reprogramming process including inflammatory (Jayawardena et al., 2012; Zhou et al., 2017), Notch (Abad et al., 2017), Wnt (Ifkovits et al., 2014), and pro-fibrotic signaling pathways (Ifkovits et al., 2014; Zhao et al., 2015). However, the mechanisms by which these diverse pathways impair cardiac reprogramming remains poorly understood.

Widespread re-patterning of epigenetic modifications including histone tail methylation and acetylation are associated with cellular reprogramming (Theunissen and Jaenisch, 2014). Previous studies have shown that histone H3 lysine 27 trimethylation (H3K27me3) is highly enriched at cardiac gene loci in non-myocytes but is removed during cardiomyogenesis (Dal-Pra et al., 2017; Liu et al., 2016; Paige et al., 2012). Moreover, previous studies demonstrated that cardiac transcription factors physically associate with H3K27me3-specific demethylases; Isl1 interacts with Jumonji Domain-Containing Protein 3 (JMJD3) (Wang et al., 2016) whereas GATA4, NKX2.5, and TBX5 interact with Ubiquitously-Transcribed X Chromosome Tetratricopeptide Repeat Protein (UTX) (Lee et al., 2012). Knockdown of either *Kdm6a* (UTX) or *Kdm6b* (JMJD3) profoundly disrupts cardiomyogenesis. *Kdm6a* knockout prevented cardiac differentiation of embryonic stem cells (ESCs) and largely disrupted cardiac looping in female embryonic mice (Lee et al. 2012). Similarly, depletion of *Kdm6b* impaired cardiac differentiation of ESCs, similar to *Isl1* knockdown (Wang et al., 2016). Furthermore, double knockdown of *Kdm6a* and *Kdm6b* modestly reduced expression of *Gata4*, *Mef2c*, and *Tbx5* in fibroblasts reprogrammed by microRNA cocktail (Dal-Pra et al., 2017). In addition to cardiac transcription factors, other transcription factors including transforming growth factor beta (TGF-β) signaling effectors SMAD2/3 are known to interact with H3K27me3 demethylases (Estaras et al., 2012). Moreover, activation of pro-fibrotic genes via TGF-β signaling impedes cardiac reprogramming (Ifkovits et al., 2014; Zhao et al., 2015). Therefore, while previous studies demonstrate the necessity of H3K27me3 demethylases in heart development, mechanisms that regulate transcription factor-mediated recruitment of these demethylases to chromatin during cardiomyogenesis remain elusive.

In addition to interacting with cardiac transcription factors, UTX was also found to bridge the interaction between cardiac transcription factors and the SWItch/Sucrose Non-Fermentable (SWI/SNF) ATP-dependent chromatin remodeling complex subunit Brahma-related gene 1 (BRG1, encoded by the *Smarca4* gene) (Lee et al., 2012). Myocardial depletion of BRG1 led to embryonic lethality and was accompanied by a thinning of the myocardium, failure to form the interventricular septum, and reduced cardiomyocyte proliferation (Hang et al., 2010). Moreover, silencing of *Smarcd3*, which encodes a heart-enriched subunit of the SWI/SNF complex, BAF60C, also led to embryonic lethality and impaired heart development and cardiomyocyte differentiation (Lickert et al., 2004).

Our previous report demonstrated that inhibition of pro-fibrotic signaling pathways including the TGF-β pathway alongside overexpression of <u>G</u>ATA4, <u>H</u>AND2, <u>M</u>EF2C, <u>T</u>BX5, miR-1, and miR-133 (GHMT2m) reprogrammed mouse embryonic fibroblasts (MEFs) with approximately 60% efficiency (Zhao et al., 2015). Here, we identify novel interactions between JMJD3 and cardiac transcription factors GATA4, HAND2, and TBX5. Depletion of JMJD3 or inhibition of its H3K27me3 demethylase activity dramatically reduces GHMT2m-mediated reprogramming efficiency. In addition, we show that the canonical TGF-β signaling effector SMAD2 disrupts the interactions between GATA4 and chromatin modifiers JMJD3 and BRG1. Furthermore, SMAD2 overexpression or TGF-β stimulation results in inefficient demethylation of H3K27me3 at cardiac genes corresponding with decreased transcription and reduced reprogramming efficiency. A point mutation in *GATA4* (c.799G>A, (Tang et al., 2006; Wang et al., 2013)) associated with congenital heart disease disrupts interaction between GATA4 and JMJD3. Human induced pluripotent stem cells (hiPSCs) carrying this mutation exhibit increased H3K27me3 levels at cardiac loci and impaired cardiogenesis. Our data establish a novel mechanism through which canonical TGF-β signaling controls cardiomyogenesis.

## Results

### Canonical TGF-β signaling effector SMAD2 impairs cardiac reprogramming

Our previous reports indicate that pro-fibrotic signaling pathways including TGF-β are potent endogenous barriers to cardiac reprogramming (Riching et al., 2018; Zhao et al., 2015). Canonical TGF-β signaling results in phosphorylation of downstream effectors SMAD2 and SMAD3, which then translocate to the nucleus via SMAD4 to activate TGF-β responsive genes. We therefore overexpressed GFP, SMAD2, or SMAD7 (an endogenous inhibitor of SMAD2 and SMAD3 phosphorylation) in GHMT2m reprogrammed cells. Overexpression of SMAD2 increased levels of phosphorylated SMAD2 by two-fold, comparable to TGF-β1 stimulation (**Figures 1A and 1B**). In contrast, SMAD7 overexpression reduced levels of phosphorylated SMAD2, as did treatment with the type I TGF-β receptor inhibitor A-83-01 (**Figure 1A and 1B**). Overexpression of SMAD7 or treatment with A-83-01 significantly increased the number of beating iCMs at days 10 and 13 whereas overexpression of SMAD2 dramatically reduced the number of beating iCMs at these time points, comparable to TGF-β1 treatment (**Figure 1C and Movies S1-5**). Interestingly, A-83-01 treatment generated the greatest number of beating cells, which may be due to the higher efficiency in blocking phosphorylation of SMAD2 than SMAD7 overexpression (**Figure 1A**). Strikingly, A-83-01 treatment of reprogramming cells overexpressing SMAD2 produced similar numbers of beating iCMs to GFP control, which was accompanied by nearly undetectable levels of phosphorylated SMAD2 levels in both conditions (**Figure S1A and S1B**), suggesting that A-83-01 enhances cardiac reprogramming through inhibiting canonical TGF-β signaling. SMAD7 overexpression resulted in two- to four-fold upregulation of cardiac genes *Myh6, Actc1,* and *Pln*, comparable to A-83-01 treatment (**Figure 1D**). In contrast, SMAD2 overexpression and TGF-β1 treatment resulted in reduced expression of these genes compared to GFP control (**Figure 1C and Movies S1-5**). Finally, SMAD7 overexpression promoted better sarcomere organization by Day 9 compared to GFP control whereas SMAD2 overexpression resulted in significantly more sarcomere disarray (**Figure 1E and 1F**).

**Figure 1:**
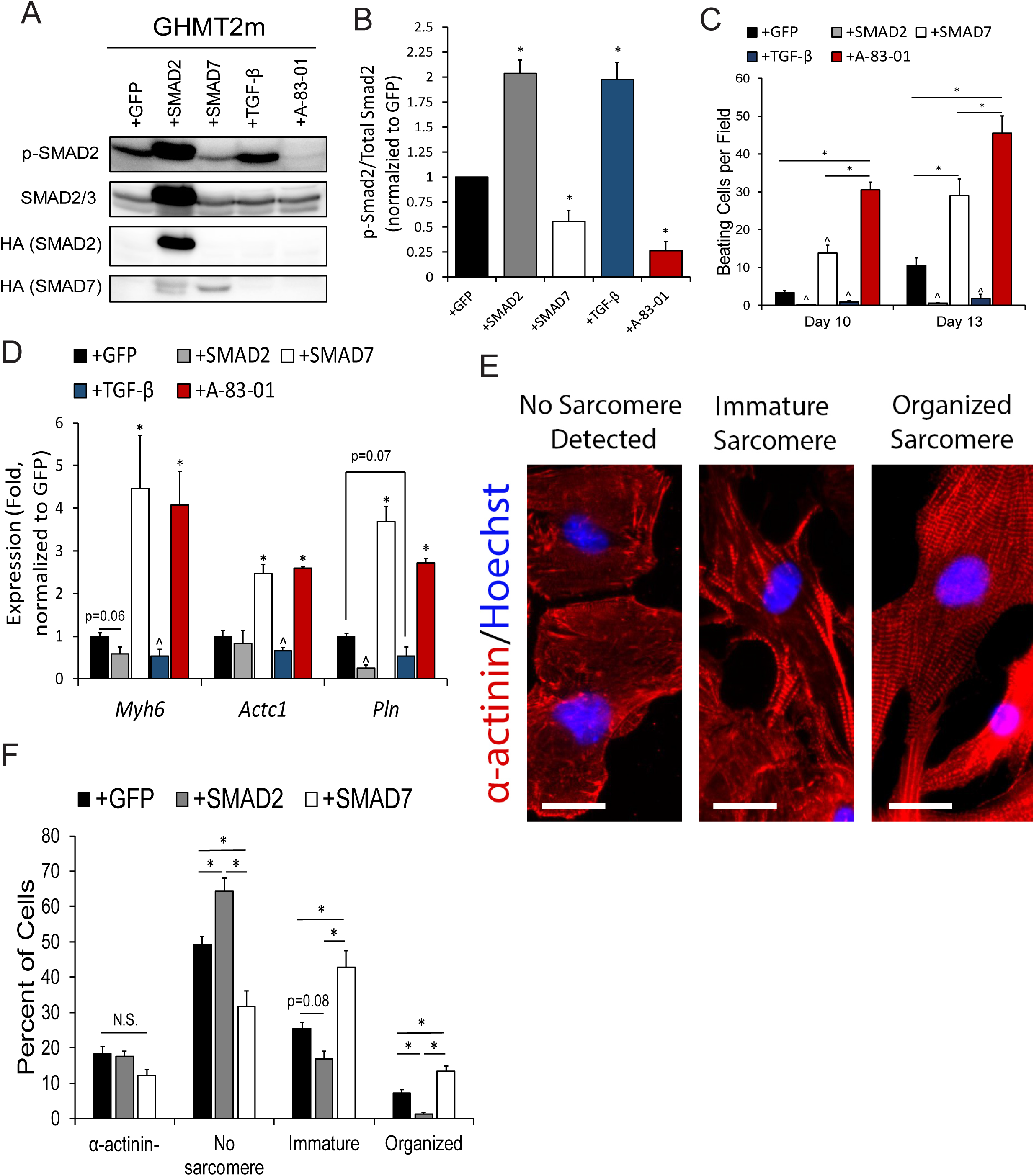
Activation of Canonical TGF-β signaling impairs cardiac reprogramming. **A and B)** Immunoblot (A) and quantification (B) of phosphorylated SMAD2 (p-SMAD2) normalized to Total SMAD2 from whole cell extracts of Day 12 GHMT2m-reprogrammed cells co-infected with GFP, SMAD2, or SMAD7, or treated with 5 ng/mL TGF-β1 or 0.5 µM A-83-01. N = 3 per group. **C)** Beating cell counts per field (0.89 mm^2^) from Day 10 and Day 13 GHMT2m-reprogrammed cells co-infected with GFP, SMAD2, or SMAD7, or treated with 5 ng/mL TGF-β1 or 0.5 µM A-83-01. N = 3 per group. **D)** Messenger RNA expression of cardiac genes *Myh6*, *Actc1*, and *Pln* harvested from Day 13 GHMT2m-reprogrammed MEFs co-infected with GFP, SMAD2, or SMAD7, or treated with 5 ng/mL TGF-β1 or 0.5 µM A-83-01. All data were normalized to the GFP group. N = 3 per group. Representative fluorescent images of indicated levels of sarcomere organization in Day 9 GHMT2m-reprogrammed MEFs. Red: α-actinin; Blue: Hoechst. Scale bar = 25 µM. **E)** Quantification of sarcomere organization in Day 9 GHMT2m-reprogrammed MEFs co-infected with GFP, SMAD2, or SMAD7. 10 fields of view per dish were collected across 3 individual experiments per group to analyze sarcomere organization. All data shown as mean ± SEM. * *p*<0.05 by one-way ANOVA with Tukey’s multiple comparison test vs the GFP group unless otherwise specified. ^ *p*<0.05 by Student’s *t* test vs the GFP group.

Taken together, these findings indicate that canonical TGF-β signaling impairs cardiac reprogramming via the effector SMAD2.

### Cardiac transcription factors physically interact with JMJD3

Removal of the transcriptionally repressive epigenetic mark H3K27me3 has been observed during cardiac reprogramming (Dal-Pra et al., 2017; Liu et al., 2016). We therefore tested the dynamics of H3K27me3 removal in our system by performing chromatin immunoprecipitation (ChIP) followed by quantitative PCR (ChIP-qPCR). In agreement with previous reports, we observed significant reduction of H3K27me3 at the *Gata4* promoter by Day 3 in GHMT reprogrammed cells compared to undifferentiated MEFs (**Figure 2A**). H3K27me3 removal proceeded through Day 7, but not to the extent observed in neonatal mouse CMs (NMCMs) (**Figure 2A**). Previous reports indicate that the H3K27me3 demethylase JMJD3 physically interacts with SMAD2/3 (Estaras et al., 2012) whereas the H3K27me3 demethylase UTX interacts with GATA4, NKX2.5, and TBX5 (Lee et al., 2012). To investigate these interactions, we overexpressed HA-tagged UTX or JMJD3 alongside MYC-tagged GHMT in HEK293T cells and performed co-immunoprecipitations (Co-IPs). In agreement with previous studies, our data confirm JMJD3 physically interacts with SMAD2/3 and UTX weakly interacts with GATA4 and TBX5 (**Figure 2B and Figure S2A**). In addition, we observed strong interaction between JMJD3 and GATA4, TBX5 and HAND2 (**Figure 2B-D**).

**Figure 2:**
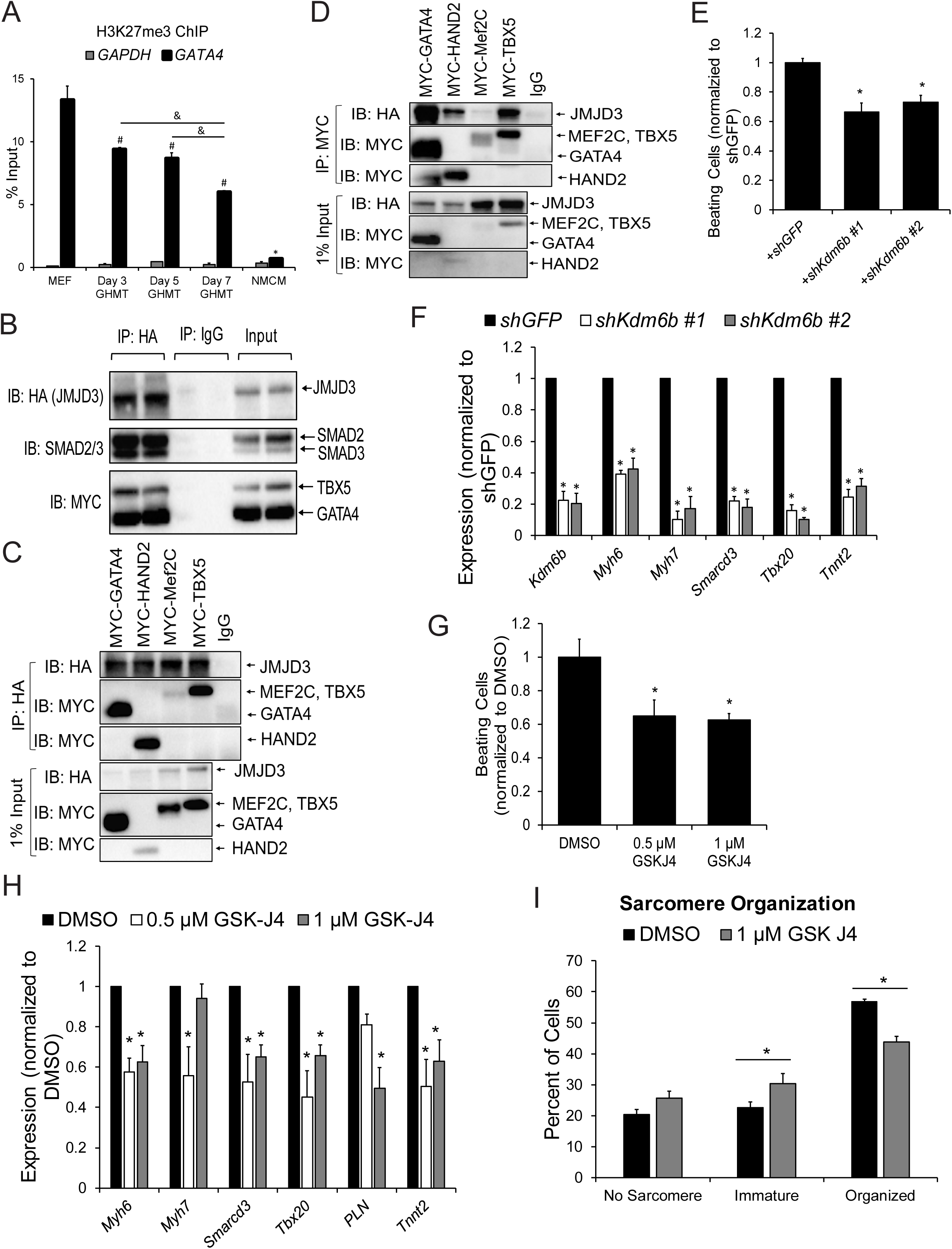
GHMT Reprogramming Factors Interact with JMJD3 to promote demethylation of H3K27me3. **A)** H3K27me3 levels in Day 3, 5, and 7 GHMT-reprogrammed cells at the *Gata4* and *Gapdh* promoters. Undifferentiated MEFs and neonatal mouse cardiomyocytes (NMCMs) were used as positive and negative controls respectively for *Gata4* H3K27me3 levels. *Gapdh* served as a negative control for H3K27me3 in all samples. All data are presented as percentage of input. *, #, and & *p*<0.05 by one-way ANOVA with Tukey’s multiple comparison test vs all groups (*), MEFs (#), or specified comparisons (&). **B)** Co-IPs using nuclear lysates harvested from HEK293T cells transfected with MYC-GHMT and HA-JMJD3. Following Co-IP with anti-HA, proteins were resolved by SDS-PAGE and immunoblotted for MYC, HA, and SMAD2/3. **C and D)** Co-IPs using nuclear lysates harvested from HEK293T cells transfected with HA-JMJD3 and individual MYC-G, -H, -M, or -T transcription factors. Following Co-IP with anti-HA (C) or anti-MYC (D), proteins were resolved by SDS-PAGE and immunoblotted for MYC and HA. **E)** Beating cell counts per field (0.89 mm^2^) from Day 12 GHMT2m-reprogrammed cells co-infected with shGFP or two different shRNA sequences targeting *Kdm6b*. Reprogrammed cells were maintained in 0.5 µM A-83-01 starting at Day 3. N = 3 per group. **F)** Messenger RNA expression of *Kdm6b* and cardiac genes *Myh6*, *Myh7*, *Smarcd3*, *Tbx20*, and *Tnnt2* harvested from Day 13 GHMT2m-reprogrammed MEFs co-infected with shGFP or two different shRNAs targeting *Kdm6b*. Reprogrammed cells were maintained in 0.5 µM A-83-01 starting at Day 3. All data were normalized to the GFP group. N = 3 per group. **G)** Beating cell counts per field (0.89 mm^2^) in Day 12 GHMT2m-reprogrammed cells treated with DMSO, 0.5 µM, or 1 µM GSK-J4. GSK-J4 was added to cells between Days 3 and 5. Reprogrammed cells were maintained in 0.5 µM A-83-01 starting at Day 3. N = 3 per group. **H)** Messenger RNA expression of cardiac genes *Myh6*, *Myh7*, *Smarcd3*, *Tbx20*, *Pln*, and *Tnnt2* harvested from Day 13 GHMT2m-reprogrammed MEFs treated with DMSO, 0.5 µM, or 1 µM GSK-J4. GSK-J4 was added to cells between Days 3 and 5. Reprogrammed cells were maintained in 0.5 µM A-83-01 starting at Day 3. All data were normalized to the GFP group. N = 3 per group. **I)** Quantification of sarcomere organization in Day 12 GHMT2m-reprogrammed MEFs treated with DMSO or 1 µM GSK-J4. GSK-J4 was administered between Days 3 and 5. Reprogrammed cells were maintained in 0.5 µM A-83-01 starting at Day 3. 10 fields of view per dish were collected across 3 individual experiments per group to analyze sarcomere organization. * *p*<0.05 by Student’s *t* test. All data shown as mean ± SEM. * *p*<0.05 by one-way ANOVA with Tukey’s multiple comparison test vs the control (shGFP or DMSO) groups.

Together, our data indicate that cardiac transcription factors GATA4, HAND2, and TBX5 as well as TGF-β signaling effectors SMAD2 and SMAD3 physically interact with the H3K27me3 demethylase JMJD3.

### JMJD3 is required for cardiac reprogramming

Our findings demonstrate JMJD3 physically interacts with cardiac transcription factors, but the role of JMJD3 in cardiac reprogramming remains unknown. Therefore, we utilized short hairpin RNA (shRNA)-mediated gene silencing to knock down JMJD3 (*Kdm6b*) expression in GHMT2m reprogrammed cells. Knockdown of *Kdm6b* using shRNAs decreased expression of *Kdm6b* by approximately 80% and decreased the number of beating cells by approximately 40%, compared to GFP shRNA control (**Figure 2E and 2F**). In contrast, cells expressing *Kdm6a* (UTX) shRNAs yielded fewer beating cells during early stages of reprogramming (Day 9) but were indistinguishable from cells expressing shGFP by Day 13 (**Figure S2B**). Moreover, *Kdm6b* knockdown in reprogrammed cells resulted in significantly reduced mRNA expression of the cardiac genes *Myh6, Myh7, Smarcd3, Tbx20,* and *Tnnt2* (**Figure 2F**). To determine whether H3K27me3 demethylase activity is essential in cardiac reprogramming, we utilized the H3K27me3 demethylase inhibitor GSK-J4 (Kruidenier et al., 2012). Treatment with 0.5 µM GSK-J4 resulted in two- to three-fold increased H3K27me3 levels at cardiac gene promoters compared to DMSO treatment (**Figure S3A**). Importantly, treatment with GSK-J4 did not interfere with the ability of GATA4 to bind target gene promoters *Tbx20, Myh6*, or *Nppa* (**Figure S3B**) or the ability of GATA4 to bind JMJD3 (**Figure S3C**). Treatment of GHMT2m reprogrammed cells with 0.5 or 1 µM GSK-J4 yielded approximately 40% fewer beating cells compared to DMSO treatment (**Figure 2G**), similar to *Kdm6b* knockdown. Additionally, treatment of GHMT2m reprogrammed cells with 0.5 or 1 µM GSK-J4 significantly reduced expression of *Myh6, Smarcd3, Tbx20*, and *Tnnt2* but only 1 µM GSK-J4 significantly reduced expression of *Pln* (**Figure 2H**). Finally, GHMT2m reprogrammed cells treated with 1 µM GSK-J4 resulted in increased sarcomere disarray compared to DMSO control (**Figure 2I**). These results demonstrate that the H3K27me3 demethylase activity of JMJD3 but not UTX is essential for cardiac reprogramming.

### Activated TGF-β signaling prevents demethylation of H3K27me3 and reduces the binding of GATA4 to target genes

Our data indicate GATA4, HAND2, TBX5, SMAD2, and SMAD3 physically interact with JMJD3 (**Figure 2B-D**) and that the H3K27me3 demethylase activity of JMJD3 is required for cardiac reprogramming. Therefore, we inquired whether TGF-β pathway activation alters the H3K27me3 landscape during reprogramming. We profiled the global H3K27me3 landscape by performing H3K27me3 ChIP followed by massively parallel Next Generation sequencing (ChIP-seq) in Day 7 GHMT2m reprogrammed cells and undifferentiated MEFs (**Figure 3A**). We manipulated TGF-β signaling in reprogrammed cells by either overexpressing SMAD2 or treating with 0.5 µM A-83-01 as these conditions led to the greatest changes in phosphorylation of SMAD2 and numbers of beating cells (**Figure 1A and 1C**). All reprogramming conditions exhibited reduced H3K27me3 levels at cardiogenic promoters including *Gata4* and *Tbx20* compared to undifferentiated MEFs (**Figure 3B-C**). Our previous RNA-seq analysis identified 509 genes upregulated >two-fold in reprogrammed cells treated with A-83-01 vs those treated with DMSO which corresponded with cardiac muscle development and mitochondrial function pathways by Gene Ontology (GO) analysis (Zhao et al., 2015). We selected one of the top enriched cardiac-specific GO terms (Regulation of Heart Contraction, GO:0008016) and quantified the levels of H3K27me3 at all promoters within this pathway (**Figure S4A and S4B**). All reprogrammed cells exhibited reduced H3K27me3 signal compared to undifferentiated MEFs. Additionally, cardiac gene promoters in reprogrammed cells overexpressing SMAD2 contained higher H3K27me3 levels than reprogrammed cells treated with DMSO or A-83-01 (**Figure S4A and S4B**). We then further analyzed the RNA-seq data (Zhao et al., 2015, deposited in the NCBI Gene Expression Omnibus, GSE71405) and selected genes upregulated >two-fold in GHMT2m reprogrammed cells treated with A-83-01 compared to undifferentiated MEFs. GO analysis revealed a number of pathways associated with muscle differentiation, ion channel physiology, metabolism, and cardiac muscle (**Figure S4C**). We evaluated the top enriched pathway by GO analysis, Muscle System Process (GO: GO:0003012). All three reprogramming conditions displayed significantly reduced H3K27me3 levels compared to undifferentiated MEFs (**Figure 3D**). Strikingly, reprogrammed cells overexpressing SMAD2 had significantly increased H3K27me3 levels at muscle-related genes compared to DMSO or A-83-01 treated reprogrammed cells (**Figure 3D**). Similar trends were observed when plotting individual candidate cardiac genes (**Figure 3E**).

**Figure 3:**
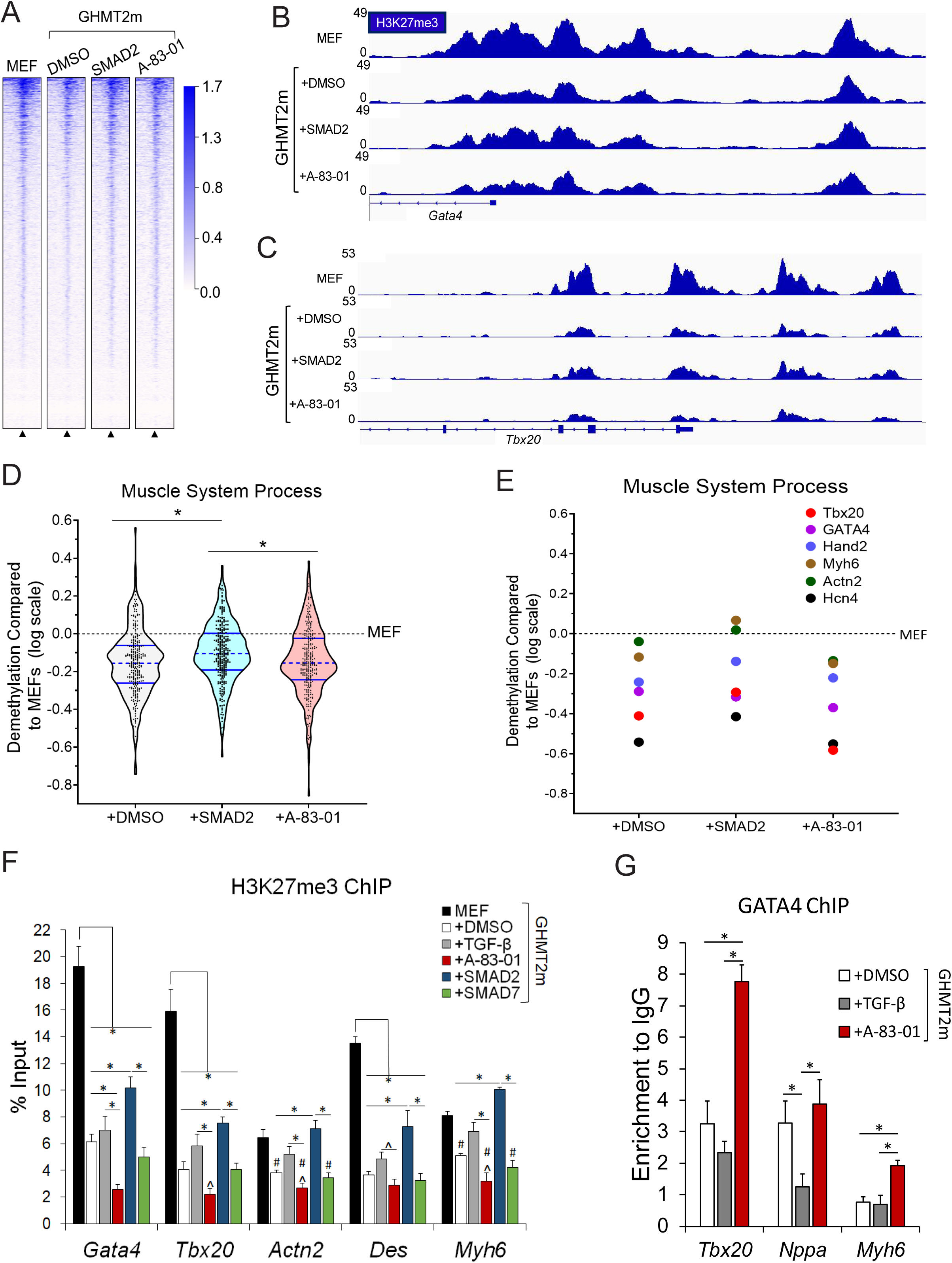
TGF-β signaling impairs demethylation of H3K27me3 and recruitment of GATA4 to target genes. **A)** Global ChIP-seq density heatmap of H3K27me3 in Day 7 GHMT2m-reprogrammed cells with indicated TGF-β pathway manipulation and undifferentiated MEFs at ± 2kb from annotated transcription start sites (TSS). **B and C)** Representative IGV tracks depicting H3K27me3 intensity at *Gata4* (B) and *Tbx20* (C) promoters from Day 7 GHMT2m-reprogrammed cells with indicated TGF-β pathway manipulation and undifferentiated MEFs. **D and E)** Logarithmic transformation of H3K27me3 levels in relation to undifferentiated MEFs at ± 2kb from the annotated TSS of genes within the Muscle System Process Gene Ontology pathway (GO:0003012) that were also upregulated >2 fold in GHMT2m + A-83-01 reprogrammed cells compared to undifferentiated MEFs as identified by RNA-seq (D) and at candidate cardiac genes selected from GO:0003012 (E). **F)** H3K27me3 levels at cardiac gene promoters *Gata4, Tbx20, Actn2, Des,* and *Myh6* from Day 7 GHMT2m-reprogrammed cells co-infected with GFP, SMAD2, or SMAD7 or treated with DMSO, 5 ng/mL TGF-β1, or 0.5 µM A-83-01 analyzed by ChIP-qPCR. Undifferentiated MEFs served as a positive control for H3K27me3 levels. Data are presented as percentage of input. N=4 for MEFs, N=6 for GHMT2m + GFP + DMSO, N=6 for GHMT2m + GFP + TGF-β1, N=5 for GHMT2m + GFP + A-83-01, N=4 for GHMT2m + SMAD2, and N=5 for GHMT2m + SMAD7. **G)** GATA4 levels at cardiac gene promoters *Nppa, Tbx20,* and *Myh6* from Day 5 GHMT2m-reprogrammed cells treated with DMSO, 5 ng/mL TGF-β1, or 0.5 µM A-83-01 analyzed by ChIP-qPCR. Data are presented as enrichment to IgG control. N=5 for GHMT2m + DMSO, N=4 for GHMT2m + TGF-β1, N=5 for GHMT2m + A-83-01. All data shown as mean ± SEM. *, # *p*<0.05 by one-way ANOVA with Tukey’s multiple comparison test at specified comparisons (*) or vs MEFs (#). ^ *p*<0.05 by Student’s *t* test vs DMSO.

We then performed ChIP-qPCR to validate our ChIP-seq model at candidate cardiac genes. We observed H3K27me3 demethylation in all reprogrammed cells at *Gata4, Tbx20,* and *Des* promoters (**Figure 3F**). Strikingly, reprogrammed cells treated with A-83-01 or overexpressing SMAD7 displayed lower levels of H3K27me3 at all promoters investigated aside from *Des*, indicating highly efficient removal of H3K27me3. In contrast, reprogrammed cells treated with TGF-β1 or overexpressing SMAD2 had consistently elevated levels of H3K27me3 compared to other reprogrammed conditions (**Figure 3F**). Notably, demethylation of H3K27me3 did not occur in reprogrammed cells treated with TGF-β1 or overexpressing SMAD2 at the *Actn2* and *Myh6* promoters (**Figure 3F**). Interestingly, vehicle treated reprogrammed cells had indistinguishable levels of H3K27me3 compared to A-83-01 treated reprogrammed cells by Day 9, suggesting that H3K27me3 demethylation at early stages is crucial for cardiac reprogramming (**Figure S4A**). In contrast, H3K27me3 levels in reprogrammed cells treated with TGF-β1 remained significantly elevated at Day 9 (**Figure S4A**). GATA4 and SMAD2/3/4 are known to form a complex (Belaguli et al., 2007); therefore, we hypothesized that activation of TGF-β signaling may recruit GATA4 away from cardiac genes. Indeed, we observed reduced binding of GATA4 to the *Nppa* promoter in Day 5 reprogrammed cells treated with TGF-β1 (**Figure 3G**). In contrast, reprogrammed cells treated with A-83-01 displayed significantly increased binding of GATA4 to *Tbx20* and *Myh6* promoters compared to TGF-β1 treated or vehicle treated reprogrammed cells (**Figure 3G**). Basal TGF-β signaling appeared to have intermediate GATA4 binding at these promoters; GATA4 binding to *Nppa* was indistinguishable from A-83-01 treated reprogrammed cells whereas GATA4 binding to *Tbx20* and *Myh6* was significantly lower than A-83-01 treated reprogrammed cells (**Figure 3G**).

Taken together, our data indicate that canonical activation of TGF-β signaling significantly impairs demethylation of H3K27me3 and binding of GATA4 to cardiac gene loci and correlates with decreased cardiac gene expression and reprogramming efficiency (**Figure 1**).

### Canonical TGF-β signaling disrupts the interactions between GATA4 and JMJD3/BRG1, thereby reducing gene expression

JMJD3 interacts with GATA4, TBX5 and HAND2 during cardiac reprogramming (**Figure 2B-D**). It has been shown that JMJD3 also acts as a scaffolding protein by bridging the interactions between BRG1 and t-box transcription factors including T-bet (Miller et al., 2010). Therefore, we hypothesized that JMJD3 and cardiac transcription factors could interact with other nuclear factors to form a complex and efficiently activate gene expression. To test this hypothesis, we performed unbiased mass spectrometry analysis to identify additional proteins that interact with GHMT. We transduced MEFs with GHMT, harvested nuclear extracts of reprogrammed cells at Day 5, and performed Co-IP followed by SDS-PAGE. Gels were stained with SPYRO Ruby protein gel stain and individual bands were excised for mass spectrometry. Our mass spectrometry analysis revealed that several components of the SWI/SNF chromatin remodeling complex including BRG1, BAF180, BAF170, and BAF155 co-immunoprecipitated with GHMT reprogramming factors (**Figure S5A and S5B**). We confirmed by Co-IP assays that both JMJD3 and GHMT reprogramming factors interact with BRG1 (**Figures 5A and B**).

**Figure 5:**
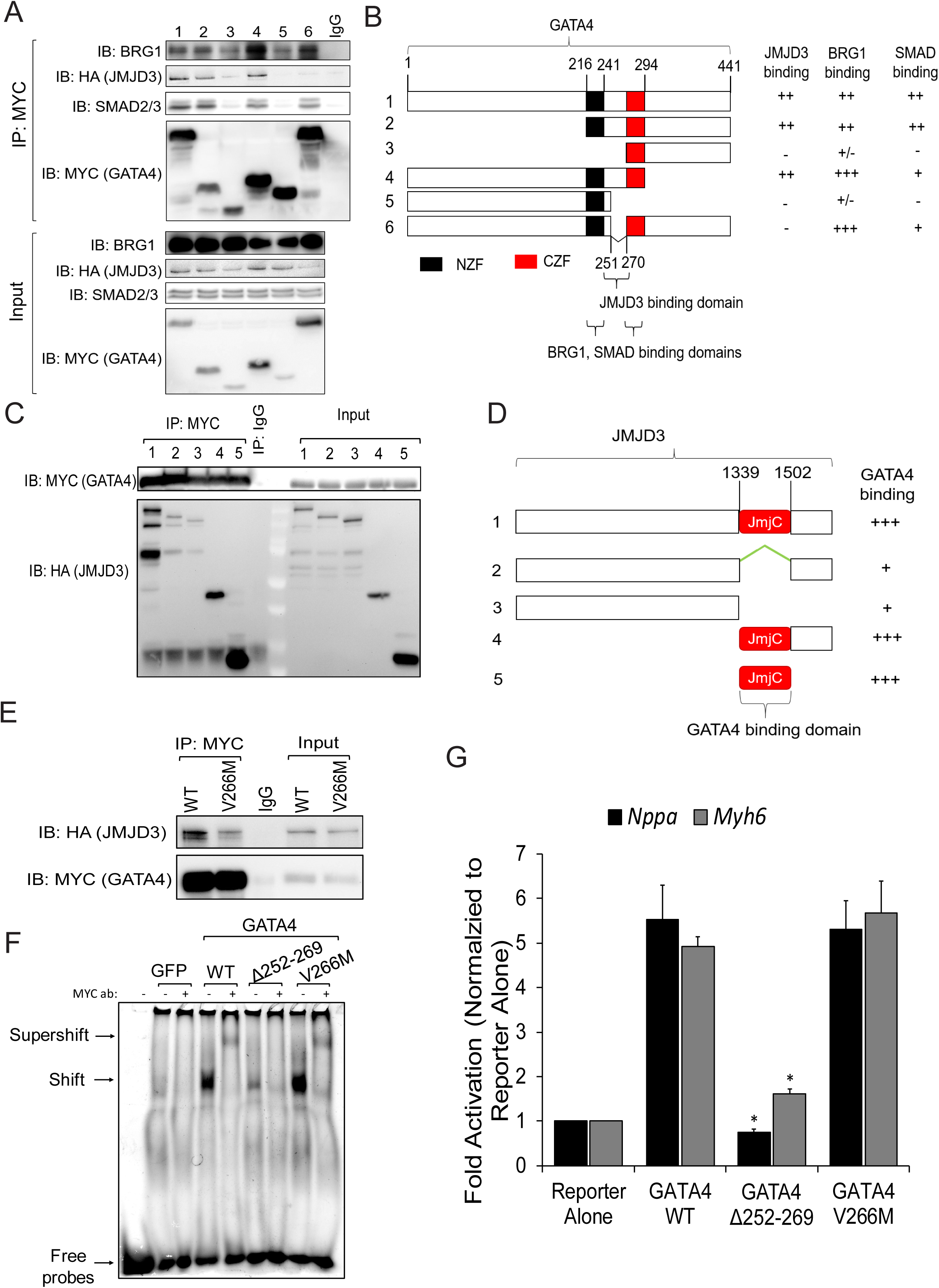
GATA4 interacts with JMJD3 via a short linker between Zinc Finger domains including residue V266. **A and B)** Co-IPs (A) using nuclear lysates harvested HEK293Ts transfected with full length HA-JMJD3 and MYC-GATA4 truncation constructs (Schematic, B). Following Co-IP with anti-MYC, proteins were resolved by SDS-PAGE and immunoblotted for BRG1, HA, SMAD2/3, and MYC. NZF, N Terminal zinc finger. CZF, C Terminal zinc finger. +++, ++, +, +/-, and – represent strong, moderate, mild, minimal, and no binding. **C and D)** Co-IP (C) using nuclear lysates harvested HEK293Ts transfected with full length MYC-GATA4 and HA-JMJD3 truncation constructs (Schematic, D). Following Co-IP with anti-MYC, proteins were resolved by SDS-PAGE and immunoblotted for HA and MYC. JmjC, Jumonji C domain. +++ and + represent strong and mild binding. **E)** Co-IPs using nuclear lysates harvested HEK293Ts transfected with HA-JMJD3 and MYC-GATA4 wildtype (WT) or MYC-GATA4 V266M mutant. Following Co-IP with anti-MYC, proteins were resolved by SDS-PAGE and immunoblotted for HA and MYC. **F)** Electrophoretic mobility shift assay (EMSA) performed using nuclear extracts from HEK293Ts transfected with GFP, WT GATA4, GATA4Δ252-269, or GATA4 V266M. Arrows point to free probes, probes bound to GATA4, or probes bound to GATA4/MYC antibody (supershift) complex. **G)** Luciferase reporter assay performed in HeLa cells co-transfected with *Nppa* or *Myh6* promoter vectors, Renilla vector, and WT GATA4, GATA4Δ252-269, or GATA4 V266M. Reporter activation was determined as a ratio of firefly luciferase to Renilla luciferase and normalized to the reporter alone group. N=3 per group. Data shown as mean ± SEM. * *p*<0.05 by one-way ANOVA with Tukey’s multiple comparison test vs WT GATA4.

Co-IP as a technique can only determine whether proteins form a complex; it does not exclude the possibility that multiple complexes are precipitated by the bait protein. To reduce the complexity of our experiments, we opted to use only a single bait protein rather than a pool of MYC-tagged bait proteins (G/H/M/T). We therefore focused on identifying which reprogramming factor was indispensable for cardiac reprogramming. To this end, we serially removed one factor from our GHMT cocktail. Whereas the majority of cells stained positive for cardiac Troponin T (cTnT) and α-actinin in GHMT-transduced MEFs, removal of any single factor significantly reduced the number of positive cells for either marker (**Figures S6A-C**). While removal of MEF2C greatly reduced the number of cells expressing α-actinin and modestly reduced the number of cells expressing cTnT, removal of GATA4 nearly abolished expression of cTnT and α-actinin in reprogrammed MEFs (**Figures S6A-C**). Since GATA4 is indispensable in cardiac reprogramming, we specifically focused on the interactions between GATA4 and chromatin modifiers. We observed that by itself, GATA4 interacts with BRG1 (**Figure 4C**) and that JMJD3 greatly enhances the binding of GATA4 to BRG1 (**Figure 4C**).

**Figure 4:**
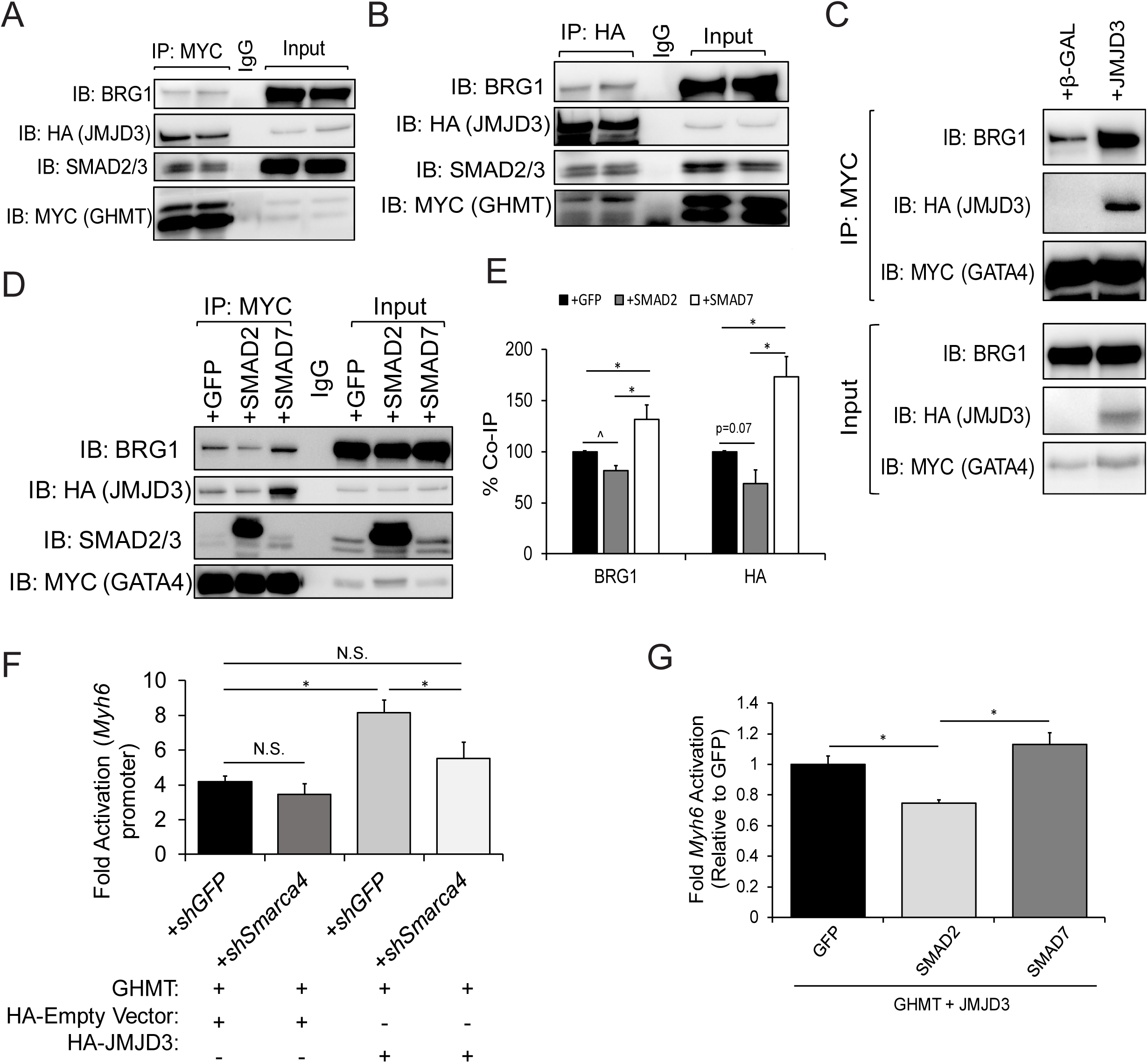
GATA4, JMJD3, and BRG1 form a large epigenetic complex which is disrupted by canonical TGF-β signaling. **A and B)** Co-IPs using nuclear lysates harvested from HEK293T cells transfected with MYC-GHMT and HA-JMJD3. Following Co-IP with anti-MYC (A) or anti-HA (B), proteins were resolved by SDS-PAGE and immunoblotted for BRG1, HA, MYC and SMAD2/3. **C)** Co-IPs using nuclear lysates harvested from HEK293T cells co-transfected with MYC-GATA4 and either β-GAL (control) or HA-JMJD3. Following Co-IP with anti-MYC, proteins were resolved by SDS-PAGE and immunoblotted for BRG1, HA, and MYC. **D)** Co-IPs using nuclear lysates harvested from HEK293Ts co-transfected with HA-JMJD3, MYC-GATA4, and GFP, SMAD2, or SMAD7. Following Co-IP with anti-MYC, lysates were resolved by SDS-PAGE and immunoblotted for BRG1, HA, SMAD2/3, and MYC. **E)** Quantification of interactions between GATA4 and JMJD3 or BRG1 shown in panel D. N=3 per group. **F)** Luciferase reporter assay performed in HEK293Ts co-transfected with *Myh6* promoter vector, Renilla vector, GHMT, pCMV-HA (empty vector control) or HA-JMJD3, and shGFP or sh*Smarca4* (BRG1). Reporter activation was determined as a ratio of firefly luciferase to Renilla luciferase and normalized to HEK293Ts transfected with *Myh6* promoter vector, Renilla vector, and pCMV-HA alone. N=3 per group. **G)** Luciferase reporter assay performed in HEK293Ts co-transfected with *Myh6* promoter vector, Renilla vector, GHMT, HA-JMJD3, and GFP, SMAD2, or SMAD7. Reporter activation was determined as a ratio of firefly luciferase to Renilla luciferase and normalized to the GFP group. N=3 per group. All data shown as mean ± SEM. * *p*<0.05 by one-way ANOVA with Tukey’s multiple comparison test at specified comparisons. ^ *p*<0.05 by Student’s *t* test.

GATA4 interacts with SMAD2/3/4 to synergistically activate gene expression in the gut epithelium (Belaguli et al., 2007). Our data indicate TGF-β signaling prevents GATA4 from binding promoters of cardiac genes (**Figure 3G**). Therefore, we hypothesized that nuclear SMAD2/3 perturbs the formation of the epigenetic complex containing GATA4, JMJD3, and BRG1. To pursue this hypothesis, we overexpressed MYC-GATA4 and HA-JMJD3 in HEK293Ts and co-expressed GFP, SMAD2, or SMAD7. Overexpression of SMAD2 significantly decreased the association between GATA4 and BRG1 (**Figure 4D and E**). While not significant, a similar trend was observed in the association between GATA4 and JMJD3 (p=0.07). In contrast, overexpression of SMAD7 enhanced the interaction between GATA4 and both BRG1 and JMJD3, indicating that canonical TGF-β signaling perturbs the formation of this epigenetic complex (**Figures 4D and E**). BRG1 has been previously shown to interact with RNA polymerase II (Zhao et al., 2005). Therefore, we hypothesized that JMJD3 promotes transcription independent of its H3K27me3 demethylase activity through its interaction with BRG1. Thus, we overexpressed GHMT and utilized an *Myh6-*luciferase reporter as a readout of transcriptional activity. Strikingly, addition of JMJD3 doubled GHMT-driven reporter activity compared to empty vector (HA) control (**Figure 4F**). Moreover, knockdown of *Smarca4* (BRG1) completely abrogated the effect of JMJD3 on reporter activity (**Figure 4F**). Finally, we assessed *Myh6*-luciferase reporter activity in response to canonical TGF-β signaling. Overexpression of SMAD2 in conjunction with GHMT and JMJD3 significantly reduced reporter activity compared to overexpression of GFP or SMAD7 (**Figure 4G**).

Altogether, our data show that GATA4 can form an epigenetic complex containing JMJD3 and BRG1 to promote transcription of cardiac genes. Additionally, the formation of this complex is disrupted by activation of canonical TGF-β signaling.

### GATA4 interacts with JMJD3 via a linker sequence between Zinc Finger domains

Our data indicate that canonical TGF-β signaling impairs cardiac reprogramming (**Figure 1**) and disrupts the interactions between GATA4 and JMJD3/BRG1 (**Figures 4D and E**). Therefore, we posited that SMAD2/3 and JMJD3 bind similar or adjacent regions of GATA4. To investigate this, we generated GATA4 truncations to map which domain(s) of GATA4 interact with BRG1, JMJD3, and SMAD2/3, respectively. Strikingly, SMAD2/3 and JMJD3 binding to GATA4 was completely abolished by removal of either zinc finger domain (**Figures 5A and B, lanes 3 and 5**). In addition, these truncation constructs also displayed reduced binding to BRG1 (**Figure 5A and B, lanes 3 and 5**). We noted that an 18 amino acid sequence (amino acids 252 to 269) residing between the two zinc finger domains of GATA4 did not overlap between truncation constructs #3 and #5 which removed the N terminal (NZF) or C terminal zinc fingers (CZF), respectively. We therefore hypothesized that this region of GATA4 was critical in binding JMJD3 and/or SMAD2/3. We generated a deletion of this 18-amino acid sequence (GATA4Δ252-269) but left both zinc finger domains intact. GATA4Δ252-269 did not exhibit impaired binding to SMAD2/3; however, binding to JMJD3 was completely abolished (**Figure 5A and B, lane 6**). Therefore, we conclude that GATA4 binds to JMJD3 via a short sequence between its two zinc finger domains, which is flanked by SMAD2/3 binding elements. We then generated truncations to map which domain(s) of JMJD3 bind GATA4. JMJD3 truncations lacking the Jumonji C (JmjC) catalytic domain exhibited reduced binding to GATA4 (**Figures 5C and D**). In contrast, the JmjC domain alone robustly co-immunoprecipitated with GATA4, suggesting the primary GATA4 binding domain resides in the JmjC domain of JMJD3 (**Figures 5C and D, lane 5**).

### Interaction between GATA4 and JMJD3 is required for GATA4 to promote cardiac reprogramming and human heart development

Our data indicate that GATA4 interacts with JMJD3 through the linker between two zinc finger domains (**Figure 5A-B**). Mutations in GATA4 are associated both with familial and sporadic cardiomyopathies (Li et al., 2014) and are also observed in patients with congenital heart disease (CHD) (Srivastava, 2006; Tomita-Mitchell et al., 2007). Mechanistically, previous studies demonstrated that the GATA4(G296S) mutation caused cardiomyopathy or congenital heart disease by impairing the ability of GATA4 to bind DNA, transcriptional cofactors, and transcriptional co-repressors, which coincided with transcriptional dysregulation and epigenetic repatterning at endothelial and cardiogenic genes (Ang et al., 2016; Garg et al., 2003). To determine whether the interaction between GATA4 and JMJD3 is crucial for GATA4 to promote cardiogenesis, we screened the Human Gene Mutation Database (http://www.hgmd.cf.ac.uk) and identified several mutations residing within the JMJD3 binding domain (amino acids 252-269). One such mutation (GATA4 c.799G>A, which encodes a missense from valine to methionine at residue 267) was associated with cardiac defects including ventricular-septal defect (VSD) and patent ductus arteriosus (PDA) (Tang et al., 2006; Wang et al., 2013). We used site-directed mutagenesis to generate a homologous mutation in mouse GATA4, GATA4(V266M). GATA4(V266M) exhibited significantly impaired binding to JMJD3 (**Figure 5E and S7**). However, this mutation did not affect the ability of GATA4 to bind DNA or activate transcription of target genes including *Myh6* and *Nppa* compared to wildtype in *in vitro* luciferase assays (**Figure 5F and G**). In contrast, GATA4Δ252-269 was unable to bind DNA, resulting in significantly impaired transcriptional activation of target genes (**Figure 5F and G**).

Since GATA4(V266M) exhibits reduced JMJD3 binding rather than altered DNA binding and/or transcriptional activity, it is an ideal mutant to determine the roles of the JMJD3-GATA4 interaction in cardiogenesis. We therefore reprogrammed MEFs with wildtype (WT), Δ252-269, or GATA4(V266M) in addition to HMT2m reprogramming factors. Since GATA4Δ252-269 is unable to bind DNA, we speculated MEFs transduced with this construct would fail to reprogram. Indeed, MEFs transduced with GATA4Δ252-269 generated almost no beating cells by Days 10 or 13 and displayed significantly reduced expression of cardiac genes at Day 5 (**Figures 6A and B**). In contrast, MEFs transduced with GATA4(V266M) did generate beating CMs, but to a lesser extent than MEFs transduced with WT GATA4 (**Figure 6A**). Moreover, expression of cardiac genes *ACTC1*, *Tbx20*, and *Smarcd3* was impaired in MEFs transduced with GATA4 V266M compared to WT GATA4 at Day 5 (**Figure 6B**).

**Figure 6:**
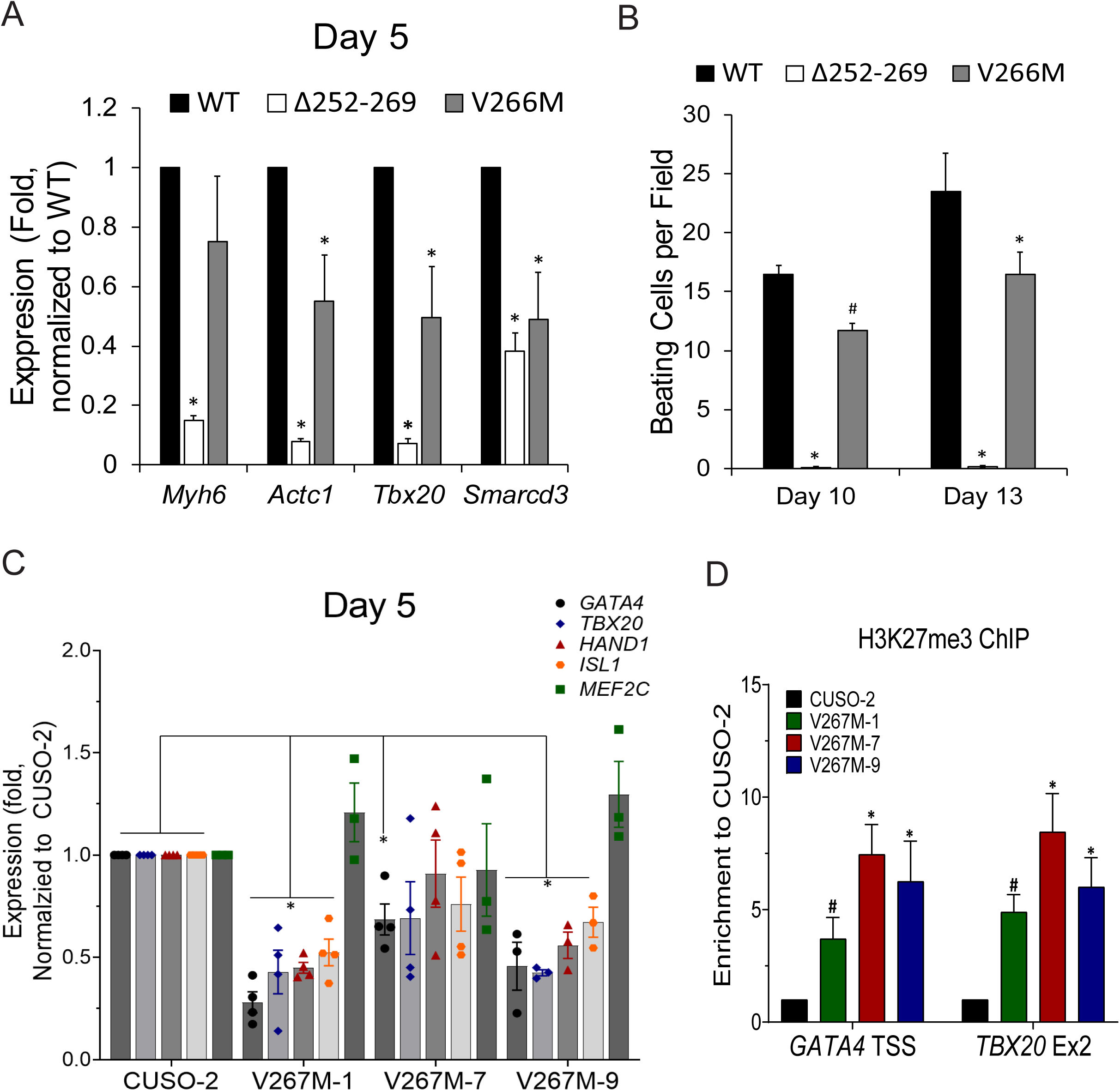
GATA4 mutation associated with CHD impairs cardiac differentiation. **A)** Messenger RNA expression of cardiac genes *Myh6*, *Actc1*, *Tbx20*, and *Smarcd3* harvested from Day 5 GHMT2m-reprogrammed using WT, Δ252-269, or V266M GATA4. Reprogrammed cells were maintained in 0.5 µM A-83-01 starting at Day 3. All data were normalized to the WT GATA4 group. N = 3 per group. **B)** Beating cell counts per field (0.89 mm^2^) from Day 10 and Day 13 GHMT2m-reprogrammed cells using WT, Δ252-269, or V266M GATA4. Reprogrammed cells were maintained in 0.5 µM A-83-01 starting at Day 3. All data were normalized to the WT GATA4 group. N = 3 per group. **C)** Messenger RNA expression of cardiac genes *GATA4*, *TBX20*, *HAND1*, *ISL1*, and *MEF2C* harvested from Day 5 WT (CUSO-2) or three isogenic V267M GATA4 hiPS-CM lines. N = 4 for CUSO-2, V267M-1, and V267M-7 lines. N= 3 for the V267M-9. **D)** H3K27me3 levels at cardiac gene promoters *Gata4* and *Tbx20* from from Day 7 WT or three isogenic V267M GATA4 hiPS-CM lines. N = 4 per group. All data shown as mean ± SEM. * *p*<0.05 by one-way ANOVA with Tukey’s multiple comparison test vs WT GATA4 groups. # *p*<0.05 by Student’s *t* test vs WT GATA4.

CHD arises from impaired heart development; therefore, we sought to study the GATA4(V267M) mutation in a human model that better reflects physiological development. We utilized CRISPR-Cas9 genome editing to generate a homozygous GATA4(V267M) mutation in human induced pluripotent stem cells (hiPSCs) (**Figure S8A and B**). We then differentiated GATA4(V267M) and WT GATA4 (CUSO-2) hiPSC lines into CMs using previously published methods (Chi et al., 2019; Lian et al., 2013). GATA4 protein levels in GATA4(V267M) lines were significantly reduced at Day 7 of differentiation (**Figure S8C**). Moreover, mRNA expression of cardiac markers *GATA4, TBX20, HAND1,* and *ISL1* was significantly reduced at Day 5 of differentiation in two GATA4 V267M lines, with similar trends in the third line (**Figure 6C**). Strikingly, H3K27me3 levels at *Gata4* and *Tbx20* promoters were markedly increased in GATA4 V267M lines compared to WT control (**Figure 6D**). Our findings reveal a novel mechanism by which mutant GATA4 causes impaired cardiac differentiation, which may translate to CHD in humans, by interfering with interactions between the transcription factor and epigenetic modifiers such as JMJD3.

## Discussion

Cardiac reprogramming represents a promising novel therapeutic approach to regenerate CMs lost after ischemic injury. While significant progress has been made in optimizing reprogramming efficiency (Zhao et al., 2015; Zhou et al., 2015; Zhou et al., 2017; Zhou et al., 2016) since the initial report (Ieda et al., 2010), mechanisms governing cardiac reprogramming remain largely unknown. Here we report a novel mechanism by which canonical TGF-β pathway activation impairs cardiac reprogramming via disrupting interactions between GATA4 and epigenetic modifiers JMJD3 and BRG1. This results in elevated levels of H3K27me3 and lower levels of GATA4 bound to cardiac promoters. Furthermore, a mutation in GATA4 (V267M), which is associated with human CHD, disrupted binding to JMJD3 and impaired demethylation of H3K27me3 at cardiac loci. This was accompanied by reduced cardiac gene expression and impaired cardiomyogenesis in hiPS-CMs carrying the mutation (GATA4 c.799G>A) or reprogrammed fibroblasts overexpressing mutant GATA4(V266M) (Figure 7). Our study therefore sheds new light on how recruitment of epigenetic complexes regulates cardiomyogenesis.

**Figure 7:**
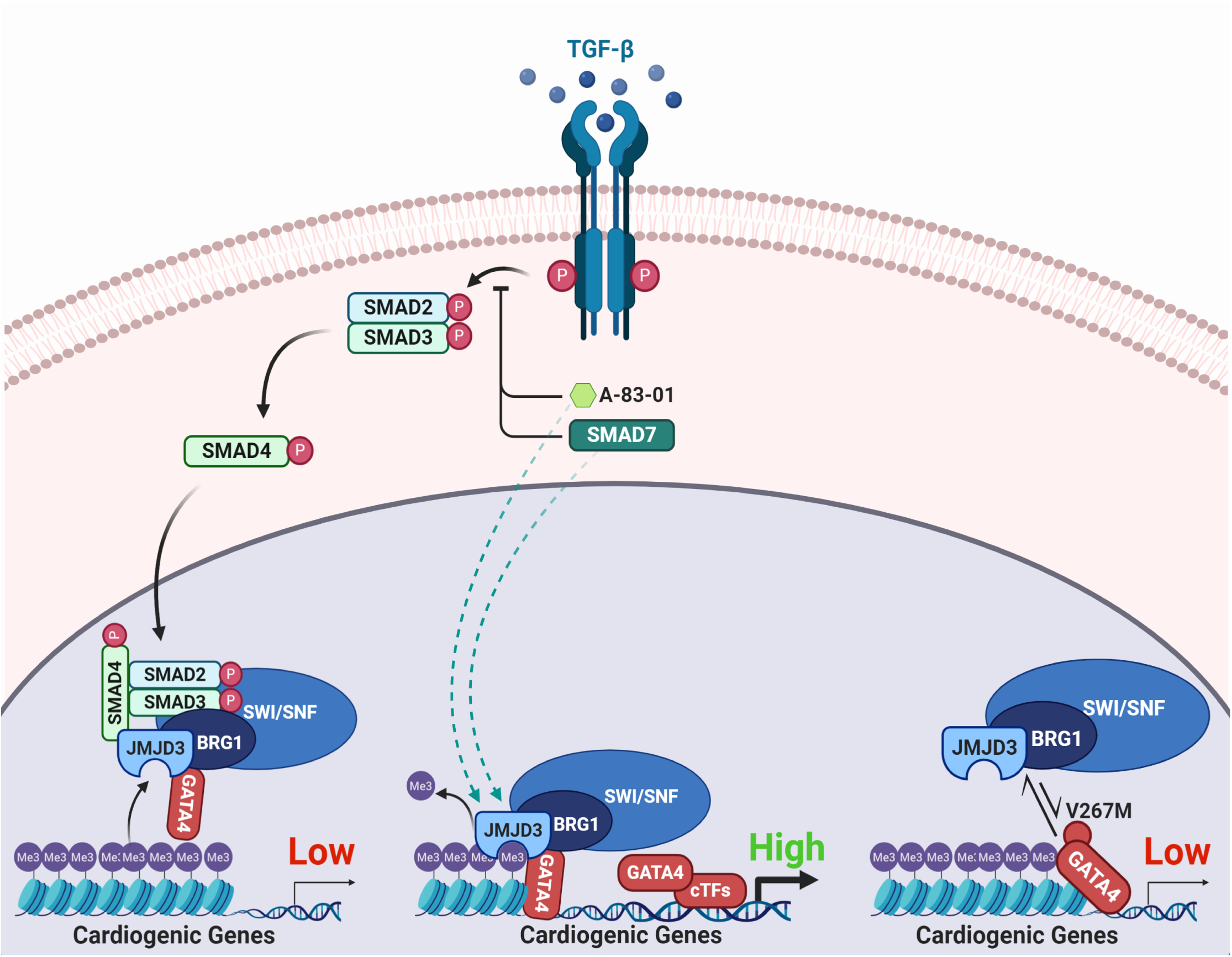
Recruitment mechanisms of epigenetic modifiers to cardiogenic genes by GATA4. Schematic summary of the regulation of interactions between GATA4 and epigenetic modifiers. Canonical TGF-β pathway activation or the congenital heart disease-associated mutation in GATA4 (V267M) impede the recruitment of epigenetic modifiers JMJD3 and BRG1 to cardiogenic genes by GATA4, resulting in impaired H3K27me3 demethylation and gene expression in cardiomyogenesis. TGF-β blockade prevents nuclear translocation of the SMAD2/3/4 complex, promoting the recruitment of epigenetic modifiers to cardiogenic genes by GATA4 resulting in efficient removal of H3K27me3 and gene expression.

TGF-β signaling results in increased deposition of H3K27me3 at genes silenced by epithelial-to-mesenchymal transition including *Cdh1* (Tange et al., 2014). Moreover, deposition of H3K27me3 was accompanied by elevated binding of Polycomb repressive complex 2 (PRC2) members EZH2 and JARID2 (Tange et al., 2014). We observed similar trends towards elevated levels of H3K27me3 at the *Myh6* promoter in GHMT2m-reprogrammed cells overexpressing SMAD2 compared to control MEFs. This suggests that canonical TGF-β signaling may also actively repress promoters, potentially via interactions with PRC2 components. Previous studies have shown that inhibition of EZH2 improves reprogramming efficiency (Dal-Pra et al., 2017; Hirai and Kikyo, 2014). Parenthetically, the EZH2 inhibitor Tazemetostat has shown promising results in clinical trials with limited toxicity (Tremblay-LeMay et al., 2018) and was granted accelerated approval status by the FDA for the treatment of epithelioid sarcoma in early 2020. Therefore, inhibition of EZH2 may be a promising alternative to TGF-β inhibition, which despite showing favorable safety profiles in patients in more recent trials, remains a safety concern given the diverse roles TGF-β signaling plays in normal physiology (Akhurst, 2017; Connolly et al., 2012; Ganesh and Massague, 2018).

Previous studies have demonstrated the importance of H3K27me3 demethylases in heart development and cardiomyogenesis (Dal-Pra et al., 2017; Lee et al., 2012; Liu et al., 2016; Wang et al., 2016). However, to our knowledge, we are the first to report a defined mechanism by which canonical TGF-β signaling dramatically alters the epigenetic landscape of GHMT2m-reprogrammed cells. JMJD3 has been shown to interact with Isl1 and is critical for cardiac differentiation of ESCs (Wang et al., 2016). GHMT-mediated reprogramming does not induce expression of *Isl1* (data not shown), so it is not likely that the JMJD3 recruitment depends on ISL1 in cardiac reprogramming. Instead, our data show that JMJD3 interacts with additional transcription factors GATA4, HAND2, and TBX5, which are downstream of ISL1 in cardiomyogenesis. These data therefore suggest that JMJD3 is critical at multiple stages of cardiac development, from early mesodermal stages through immature CMs. We show that cardiac transcription factors strongly interact with JMJD3, but minimally interact with UTX (**Figures 2B-D**, **S2A**). Consistently, knockdown of *Kdm6a* encoding UTX impaired generation of beating iCMs at early time points but beating iCM numbers were not significantly changed compared to shGFP control at later time points (**Figure S2B**). In contrast, knockdown of *Kdm6b* encoding JMJD3 significantly reduced beating iCM counts at all time points investigated. This suggests JMJD3 can compensate for UTX loss in cardiac reprogramming, but UTX cannot compensate for JMJD3 loss. Treatment of mouse ESCs with a small molecule that induces degradation of the type II TGF-β receptor was previously reported to promote cardiac differentiation at early time points (Willems et al., 2012). Since JMJD3 plays crucial roles in cardiac differentiation of ESCs (Wang et al., 2016), it is possible that the mechanism by which canonical TGF-β pathway activation disrupts recruitment of JMJD3 to cardiogenic genes also functions in physiological development in addition to fibroblast reprogramming.

Our data indicate that canonical TGF-β signaling is the primary mechanism through which TGF-β signaling impairs cardiomyogenesis. Overexpression of SMAD2 resulted in similar levels of H3K27me3 at cardiac promoters, similar expression of cardiac gene transcripts, and similar numbers of beating iCMs as treatment with TGF-β1. Moreover, overexpression of SMAD7 trended in the same direction as treatment with A-83-01 in all assays tested. However, treatment with A-83-01 still resulted in significantly increased beating iCM counts and trends in reduced H3K27me3 at cardiac promoters compared to SMAD7 overexpression. One possibility is that SMAD7 overexpression did not reduce levels of phosphorylated SMAD2 to the extent that A-83-01 treatment did (**Figure 1A**). Alternatively, it is possible that blockade of non-canonical TGF-β signaling also positively regulates cardiac reprogramming. Therefore, downstream effectors of non-canonical TGF-β signaling should be evaluated in future studies. Additionally, SMAD7 enables activation of STAT3 (Yu et al., 2017), which has previously been linked to impairing cardiac reprogramming efficiency (Jayawardena et al., 2012). Therefore, it is possible SMAD7 overexpression, while reducing detrimental effects of canonical TGF-β signaling, promotes inflammatory signaling that inhibits cardiac reprogramming efficiency, resulting in lower reprogramming efficiency compared to A-83-01 treatment.

While our ChIP-qPCR data indicate significantly reduced levels of H3K27me3 at four out of five validated promoters in A-83-01 treated reprogrammed cells compared to vehicle (DMSO), our ChIP-seq analysis did not reveal significant differences in H3K27me3 status between these two groups globally. A caveat of ChIP assays is that they only offer a snapshot of the epigenetic landscape, which is well known to dynamically change during cell differentiation. We initially chose to evaluate H3K27me3 levels in Day 7 reprogrammed cells since we observed the highest degree of demethylation by this time point in our time course assay (**Figure 2A**) and therefore rationalized that we would see the largest differences between reprogrammed cells with activated TGF-β signaling and those with inhibited TGF-β signaling. We also noted that vehicle treated cells exhibited demethylation of H3K27 similar to A-83-01 treated cells by Day 9, suggesting that basal TGF-β signaling does not prevent but does slow the kinetics of H3K27 demethylation. Therefore, it is possible that larger differences in H3K27me3 levels between A-83-01 treated cells and vehicle treated cells would be observed at earlier time points than Day 7. Alternatively, it is possible that the reprogramming processes is sensitive to the expression of a handful of cardiogenic genes. We observed that H3K27me3 levels at the *Tbx20* promoter were significantly reduced in A-83-01 reprogrammed cells compared to vehicle treated cells (**Figures 3E and 3F**). Finally, removal of a repressive epigenetic mark like H3K27me3 does not necessarily guarantee expression of a gene. We did not assess deposition of active chromatin marks including H3K4me1, H3K4me3, or H3K27ac in this study. It is possible that A-83-01 treatment promotes deposition of these active marks, further enhancing reprogramming efficiency compared to vehicle treatment. Future studies should therefore evaluate the dynamics of the epigenetic landscape beyond H3K27me3 in the context of cardiac reprogramming.

A previous report indicated that GATA4 (V267M) displayed reduced binding to the mediator complex and BRG1, which significantly reduced its transcriptional activation of genes in hepatocellular carcinoma cells (Enane et al., 2017). Our data indicate that the mouse analog, GATA4(V266M), does not disrupt DNA binding or transcriptional activity but does exhibit reduced binding to JMJD3. This reduced interaction with JMJD3 correlated with impaired fibroblast reprogramming to CMs and CM differentiation of hiPSCs. It is possible that in hepatocytes, GATA4(V267M) cannot bind a critical cofactor and fails to recruit the mediator complex whereas in cardiac progenitor cells, this mutation can still bind cofactors that recruit the mediator complex. However, in cardiac progenitor cells, GATA4(V267M) does not efficiently recruit JMJD3 to cardiogenic genes, resulting in impaired H3K27me3 demethylation and gene expression, leading to CHD-associated phenotypes. Unraveling the myriad of protein-protein interactions involved in transcriptional activation in future studies will therefore further elucidate the specific contexts in which the same mutation can lead to different pathogenic mechanisms in different tissues.

To summarize, we have established a mechanism whereby the canonical TGF-β signaling effector SMAD2 disrupts interactions between GATA4 and epigenetic modifiers JMJD3 and BRG1, resulting in elevated levels of H3K27me3 at cardiac promoters, reduced cardiac gene transcription, and impaired binding of GATA4 to target genes. Interactions between GATA4 and these epigenetic modifiers is not only required for optimal cardiac reprogramming, but also normal heart development in human.

## Acknowledgements

The authors thank the University of Colorado School of Medicine Biological Mass Spectrometry Core Facility and the Genomics and Microarray Core Facility. ASR was supported by predoctoral fellowships from the University of Colorado Consortium for Fibrosis Research & Translation, Colorado Clinical & Translational Sciences Institute (TL1 TR001081), and the American Heart Association (18PRE34030030). RAB received a postdoctoral fellowship award from the Canadian Institutes of Health Research (FRN-216927). K.S. was supported by funds from the Boettcher Foundation, American Heart Association (13SDG17400031), University of Colorado Department of Medicine Outstanding Early Career Scholar Program, Gates Frontiers Fund, and NIH R01HL133230. We thank Dr. Aaron Johnson and Dr. David Port for insightful discussion and Dr. Jennifer Major for critical reading and editing of the manuscript. Figure 7 was created using Biorender.com.

## Author Contributions

K.S. and A.S.R. designed the experiments. A.S.R., Y.Z., Y.C., C.C., R.A.B., B.J.K., performed experiments and analyzed data. H.X. and E.D. aligned and analyzed ChIP-seq data. T.A.M., T.G.K., and P.M.B. contributed scientific discussion. A.S.R., E.D., Y.Z., Y.C., C.C., R.A.B., B.J.K., H.X., T.G.K., T.A.M., P.M.B., and K.S. prepared the manuscript.

## Declaration of Interests

The authors declare no competing interests.

## Methods

### Isolation of Mouse Embryonic Fibroblasts

Embryonic fibroblasts were isolated as described previously (Riching et al., 2018). Briefly, pregnant C57BL/6 were purchased at E13 from Charles River Laboratories. Upon arrival, animals were sacrificed and embryos at E14.5 were harvested. Upper body, head, and internal organs were discarded. The body below the liver was minced into fine pieces and digested in 0.25% Trypsin/EDTA (Gibco) for 40 minutes at 37 °C with 5% CO2. Cells were resuspended in 25 mL DMEM/High Glucose (HyClone) containing 10% fetal bovine serum (FBS; Gemini), 1.1% Penicillin-Streptomycin (Gibco), and 1.1% GlutaMAX supplement (Gibco) and plated in a 15-cm dish. Medium was changed after 24 hours. After 72 hours, MEFs were collected and frozen for future use.

### Plasmids and Cloning

pBabe-X GATA4, Hand2, Mef2c, Tbx5, miR-1, and miR-133 were generated previously (Song et al., 2012; Zhao et al., 2015). pCMV-HA-JMJD3 was a gift from Kristian Helin (Addgene plasmid #24167 (Agger et al., 2007)). pBabe-X-HA-UTX was subcloned from pCMV-HA-UTX, a gift from Kristian Helin (Addgene plasmid #24168 (Agger et al., 2007)). pBabe-X-HA-SMAD2 was subcloned from pCMV5B-HA-SMAD2, a gift from Jeff Wrana (Addgene plasmid #11734 (Eppert et al., 1996)). pBabe-puro-SMAD7-HA was a gift from Sam Thiagalingam (Addgene plasmid #37044 (Papageorgis et al., 2010)). *Kdm6a, Kdm6b,* and *Smarca4* shRNA sequences (Table S1) were designed using the online webtool http://cancan.cshl.edu/RNAi_central/RNAi.cgi?type=shRNA and oligonucleotides were purchased from IDT, amplified by PCR, and ligated into a modified pShagMagic2 miR-based vector, MSCV-PM (Addgene plasmid #27021 (Nguyen et al., 2009)). Site directed mutagenesis to generate GATA4 V266M was performed with the Q5 Site-Directed Mutagenesis Kit (NEB) as per the manufacturer’s instructions. Mutagenesis primers were designed using the NEBaseChanger web tool (http://nebasechanger.neb.com/). GATA4 and JMJD3 truncation plasmids were generated by serially removing domains from N or C termini by PCR. Primers used for site directed mutagenesis and domain truncation and shRNA sequences are listed in Table S2.

### Generation of Retrovirus and Transduction of MEFs

Retroviral infection of MEFs was performed as previously described (Riching et al., 2018). Approximately 5 x 10^6^ Platinum E cells (PE cells; Cell Biolabs) were plated in 10-cm dishes. 20 hours later, cells were transfected with 36 µL FuGENE 6 (Promega) and 12 µg of retroviral plasmid DNA. On the day of transfection, MEFs were thawed and plated into 60-mm dishes pre-coated with SureCoat (Cellutron). MEFs were allowed to settle overnight. Viral supernatant was collected from transfected PE cells at 24 and 48 hours, and filtered through a 0.45-μm cellulose filter. Polybrene (Sigma) was added to viral supernatant at a concentration of 6 μg/mL and added to dishes containing MEFs. Following the first collection of viral supernatant, fresh medium (DMEM/High Glucose (HyClone) containing 10% FBS (Gemini), 1.1% Penicillin-Streptomycin (Gibco), and 1.1% GlutaMAX supplement (Gibco) was added to PE cells. Infected MEFs were maintained in induction medium composed of DMEM/High Glucose:Medium 199 (4:1, HyClone:Gibco), 10% FBS (Gemini), 5% horse serum (Gemini), antibiotics (Gibco), 1X non-essential amino acids (Gibco), 1X essential amino acids (Gibco), 1X B-27 (Gibco), 1X insulin– selenium–transferin (Gibco), 1X MEM vitamin solution (Gibco) and 1X sodium pyruvate (Gibco) starting at 72 hours post-infection. DMSO (Thermo Scientific), 5 ng/mL TGF-β1 (Novoprotein CA59), 0.5 μM A-83-01 (Tocris 2939), or 0.5 μM or 1 μM GSK-J4 (Tocris 4594) were administered starting at 72 hours post-infection. Induction medium was changed every 2 days.

### RNA extraction and quantitative, real time PCR

Total RNA was extracted from D13 reprogrammed MEFs using TRIzol reagent (Invitrogen) and chloroform. The aqueous layer containing RNA was transferred to a new tube and RNA was precipitated using isopropanol. RNA was washed with 70% ethanol and dried. cDNA was synthesized using the iScript Reverse Transcription Supermix (Biorad). Diluted cDNA was amplified using SYBR PowerUP master mix (Applied Biosystems) and expression was determined by StepOne Real-Time PCR System (Applied Biosystems). qPCR primer sequences are listed in Table S3.

### Immunocytochemistry

MEFs were seeded and reprogrammed by GHMT or GHMT2m in 12 well plates. On days 9 and 12, cells were rinsed 1X in ice-cold PBS and fixed in 2% paraformaldehyde for 10 minutes at room temperature. Cells were washed 3X in PBS and permeabilized with 0.2% Triton X-100 for 15 minutes at room temperature. Cells were then blocked in 10% horse serum (Gemini). Primary antibodies (Cardiac Troponin T; Thermo Scientific ms-295-p - 1:400; α-actinin; Sigma A7811L - 1:400) were diluted in 10% horse serum (Gemini) and added to fixed cells for 1 hour at room temperature. Cells were washed 3X in PBS and then incubated with diluted secondary antibodies (anti-mouse Alexa 555; Life Technologies A-21422 - 1:800) and Hoechst (Life Technologies 62249 - 1:10,000) for 1 hour at room temperature in the dark. Cells were then washed 3X in PBS and imaged an EVOS FL Color Imaging System (Life Technologies).

### Co-Immunoprecipitation

HEK293T cells were split into 10 cm dishes and 12 µg DNA was transfected into cells using PEI at a 3:1 ratio of PEI:DNA. 48 hours post-transfection, nuclear lysates were prepared using the NE-PER Nuclear and Cytosolic Extraction Kit (Thermo 78835). Protein concentration was determined by BCA assay and 250 µg protein lysates were incubated with anti-HA (5 µg, UBPBio Y1071), anti-Myc (2.5 µg, BD Pharmingen 551102), or anti-IgG (5 µg, Santa Cruz sc-2025) and 20 µL Protein G DynaBeads slurry (Invitrogen 10004D) overnight at 4 °C in an end-over-end mixer. The following morning, beads were washed 4 times in wash buffer (150 mM NaCl, 50 mM Tris-Cl pH 7.4, 1 mM EDTA, 1% Triton) and boiled in 1X loading dye (75mM Tris-HCl, pH 6.8, 2.4% SDS, 12% glycerol, 0.12M DTT, 0.012% bromophenol blue). Lysates were run on a 10% polyacrylamide gel and transferred to a PVDF membrane (110mA, 16 hours, 4 °C).

### Western Blot

Cells were washed with ice-cold DPBS (Gibco) twice. Whole cell extracts were harvested with ice-cold lysis buffer (150 mM NaCl, 50 mM Tris-Cl pH 7.4, 1 mM EDTA, 1% Triton, and freshly added protease inhibitors (Complete mini tablet (Roche) and 1 mM phenylmethylsulphonyl fluoride (PMSF)). Protein concentration was determined by BCA assay and 30 µg protein lysates were loaded and resolved on a 10% polyacrylamide gel and transferred to a PVDF membrane (110mA, 16 hours, 4 °C). The following primary antibodies were used: anti-p-Smad2 (Cell Signaling 3108S, 1:1000), anti-total Smad2/3 (Cell Signaling 8685S, 1:1000), anti-HA (Rockland 600-401-384, 1:5000), anti-Myc (Santa Cruz sc-789, 1:500), anti-Brg1 (Santa Cruz sc-17796, 1:500), and anti-GAPDH (Life Technologies AM4300, 1:5000). Secondary antibodies used include the following: Goat Anti-Mouse IgG (H+L; Southern Biotech, 1031-05, 1:2,000) and Goat Anti-Rabbit IgG (Life Technologies, 65–6120, 1:2,000). Band intensities were quantified by densitometry using ImageJ software.

### Chromatin Immunoprecipitation

Chromatin from D5 control MEFs or D7 reprogrammed MEFs was isolated using the Magna ChIP A/G Chromatin Immunoprecipitation kit (Millipore 17-10085) as per the manufacturer’s instructions. Briefly, 10 cm dishes of cells were crosslinked with 1% paraformaldehyde for 9 minutes at room temperature. Cells were then quenched with glycine (125 mM final concentration) for 5 minutes at room temperature. Chromatin was sheared to approximately 300 bp fragments using a Diagenode Bioruptor 300 ultrasonicator (D5 control MEFs: 15 cycles, 30 seconds on/90 seconds off, high intensity; D7 reprogrammed MEFs: 25 cycles, 30 seconds on/30 seconds off, high intensity). Chromatin was diluted 10 times and incubated overnight at 4 °C with 20 µL Protein A/G MagnaChIP beads (Millipore) and 8 µg anti-H3K27me3 (Millipore, 07-449), 10 µg anti-GATA4 (Santa Cruz sc-1237x) or 10 µg control IgG (Millipore, PP64B) antibodies in an end-over-end mixer. The next day, immunoprecipitated chromatin was washed, eluted, and reverse crosslinked with Proteinase K for 2 hours at 62 °C. For qPCR, ChIP DNA was amplified with SYBR PowerUP master mix (Applied Biosystems). Results are expressed as percentage of input. ChIP-qPCR primer sequences are listed in Table S4. For ChIP-seq, libraries were prepared from 1 ng ChIP DNA using the Ovation Ultralow Library System V2 1-16 (NuGEN). Libraries were sequenced using the NovaSEQ 6000 platform (Illumina). To remove Illumina adapters and quality-trim the read ends, reads were filtered using BBDuk (http://jgi.doe.gov/data-and-tools/bb-tools). Bowtie2 (v. 2.3.2) was used to align the 150-bp paired-end sequencing reads to the mm10 reference human genome. Samtools (v.1.5) was used to remove unmapped reads and to randomly extract the same number of reads for all samples. Peaks were called using MACS2 (v2.1.1.20160309) (Zhang et al., 2008) with default parameters. ngs.plot.r (Shen et al., 2014) was used for generating the heatmaps and average profiles related to the TSS. Peak locations were further annotated using the ChIPseeker R package (Yu et al., 2015).

### Luciferase Reporter Assay

HEK293T cells were plated into 6-well plates and transfected with PEI and two micrograms of DNA. pGL3 backbones containing a 2kb promoter upstream of the *Myh6* gene or a 640 bp promoter upstream of the *Nppa* gene were used to assess promoter activity. 48 hours post-transfection, cells were rinsed twice in 1X PBS (Gibco) and lysed with 1X passive lysis buffer (Promega). Cells were snap-frozen in liquid nitrogen to facilitate lysis. 20 µL lysates were loaded into a 96 well plate and luminescence following administration of 50 µL Luciferase Assay Reagent II (Promega) and 50 µL Stop & Glo Reagent (Promega) was measured on a plate reader (BioTek). Firefly luciferase activity was normalized to Renilla luciferase activity and all data are normalized to pGL3-reporter construct alone.

### Mass Spectrometry Identification of GHMT-interaction partners

Nuclear lysates from D5 GHMT2m reprogrammed MEFs were prepared using the NE-PER Nuclear and Cytosolic Extraction Kit (Thermo 78835). Protein concentration was determined by BCA assay and 250 µg protein lysates were incubated with anti-Myc (2.5 µg, BD Pharmingen 551102) or anti-IgG (2.5 µg, Santa Cruz sc-2025) and 20 µL Protein G DynaBeads slurry (Invitrogen 10004D) overnight at 4 °C in an end-over-end mixer. The following morning, beads were washed 4 times in wash buffer (150 mM NaCl, 50 mM Tris-Cl pH 7.4, 1 mM EDTA, 1% Triton) and boiled in 1X loading dye (75mM Tris-HCl, pH 6.8, 2.4% SDS, 12% glycerol, 0.12M DTT, 0.012% bromophenol blue). Lysates were run on a 10% polyacrylamide gel and stained with SPYRO Ruby Protein Gel Stain (Thermo). Gels were imaged under UV light with a FluorChem 8900 imaging system (Alpha Innotech). Individual bands were excised from the gel with a sterile razor blade. Excised bands were reduced, alkylated, and digested with trypsin. Cleaved peptides were analyzed by nanoflow LC-ESI-MS/MS (Thermo Scientific LTQ Orbitrap Velos Pro). Peptide sequences were mapped using ProteinProspector and Mascot software packages.

### Electrophoretic Mobility Shift Assay (EMSA)

Nuclear lysates were prepared from HEK293T cells transfected with GFP, wildtype GATA4, GATA4Δ252-269, or GATA4 V266M using the NE-PER Nuclear and Cytosolic Extraction Kit (Thermo 78835). Protein concentration was determined by BCA assay and 10 µg protein was incubated with 1.25 µM annealed oligonucleotides (5’-TCGAGGTAATTAACTGATAATGGTGC-3’ (Garg et al., 2003; McFadden et al., 2000)) and 5X binding buffer (20mM HEPES, pH 7.5, 60 mM KCl, 0.5mM DTT, 1mM MgCl2, 4.8% glycerol, and 1.5 mg/mL BSA) for 30 minutes at room temperature. For supershift assays, 0.375 µg anti-Myc (BD Pharmingen 551102) was also added to the reaction mixture. Reactions were separated on a 5% non-denaturing polyacrylamide gel, stained with 1:10,000 SYBR Gold nucleic acid gel stain (Invitrogen S11494) for 40 minutes, and imaged under 302 nm UV light with a FluorChem 8900 imaging system (Alpha Innotech).

### CRISPR-Cas9 Genome Editing

The GATA4 V267M Alt-R CRISPR-Cas9 crRNA targeting sequence was designed using the online tool at http://crispr.mit.edu and purchased from IDT. crRNA was mixed with Alt-R CRISPR-Cas9 tracrRNA (IDT) and annealed at 95 °C for 5 minutes before cooling to room temperature. Annealed gRNA complex was added to Alt-R S.p. HiFi Cas9 Nuclease, V3 (IDT) to form the RNP complex and incubated at room temperature for 20 minutes. RNP complex was added to ssDNA GATA4 V267M repair template (IDT) and delivered to wildtype iPSCs by electroporation with the Amaxa human stem cell nucleofector starter kit (Lonza). Following electroporation, single cell clones were selected and expanded for genotyping by PCR. Genomic GATA4 V267M edits were confirmed by Sanger sequencing. GATA4 V267M crRNA, ssDNA repair template, PCR genotyping, and sequencing oligonucleotide sequences are listed in Table S2.

### iPSC culture and iPS differentiation

Cardiac differentiation of hiPSCs was performed as previously described (Chi et al., 2019; Lian et al., 2013). Briefly, hiPSCs were cultured until ∼85% confluence in mTeSR1 Plus medium (STEMCELL Technologies). Cells were dissociated with Accumax (Innovative Cell Technologies) and seeded into 24 well plates pre-coated with Matrigel (Corning) at a density of 300,000 cells per well in mTeSR1 medium containing 10 µM Y27632 (APExBIO) (Day -4). mTeSR1 medium was replenished daily for 3 additional days. At Day 0, medium was changed to RPMI 1640 (Life Technologies) containing B-27 supplement minus insulin (Life Technologies) and cells were treated with 8 µM ChIR99021 (Cayman). After 24 hours (Day 1), medium was changed to RPMI 1640 containing B-27 supplement minus insulin. After 72 hours (Day 3), 0.5 mL fresh RPMI 1640 containing B-27 supplement minus insulin was added to 0.5 mL conditioned (old) medium and added to cells along with 5 µM IWP2 (Tocris). Medium was aspirated at Day 5 and replenished with fresh RPMI 1640 containing B-27 supplement minus insulin. WT and three GATA4 V267M mutant lines were cultured and harvested at Day 5 for RNA extraction, Day 7 for protein extraction, and Day 7 for H3K27me3 ChIP.

### Statistical Analysis

Unless otherwise stated, statistical analysis was performed using ANOVA followed by Tukey post hoc test for multiple comparisons. If only two groups were compared, Student’s T test (2 tailed, unequal variance) was performed instead. Data were regarded as significant at p < 0.05. All graphs are displayed as mean ± SEM.

