## supplemental materials for "Suppression of canonical TGF-β signaling enables GATA4 to interact with H3K27me3 demethylase JMJD3 to promote cardiomyogenesis"

Figure S1

A

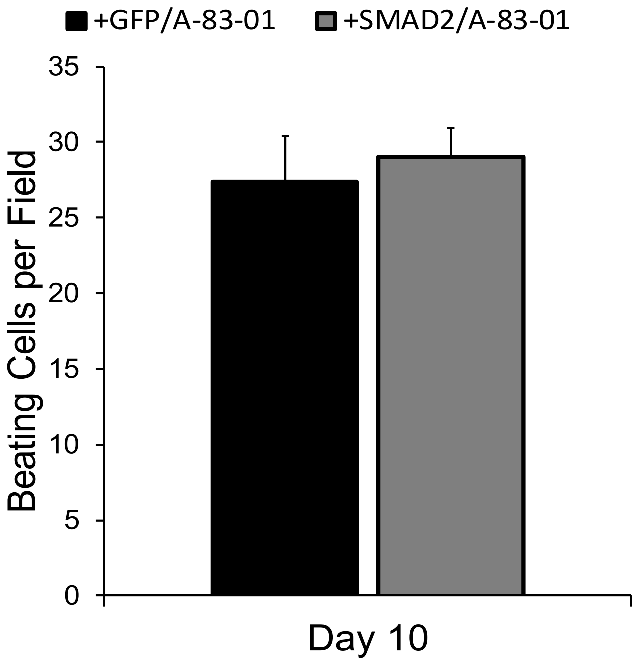

+GFP/A-83-01
+SMAD2/A-83-01
Beating Cells per Field
35
30
25
20
15
10
5
0
Day 10


B

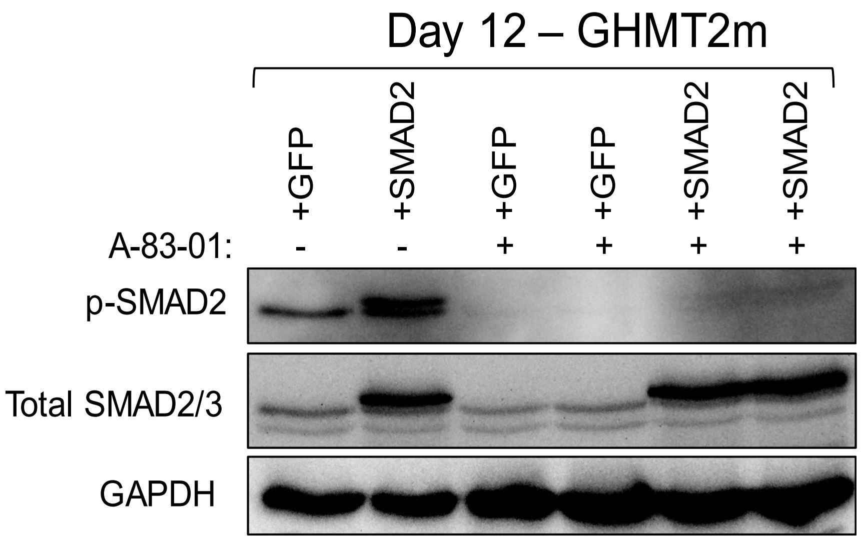

Day 12 – GHMT2m
+GFP
+SMAD2
+GFP
+GFP
+SMAD2
+SMAD2
A-83-01:
-
-
+
+
+
+
p-SMAD2
Total SMAD2/3
GAPDH

Figure S2

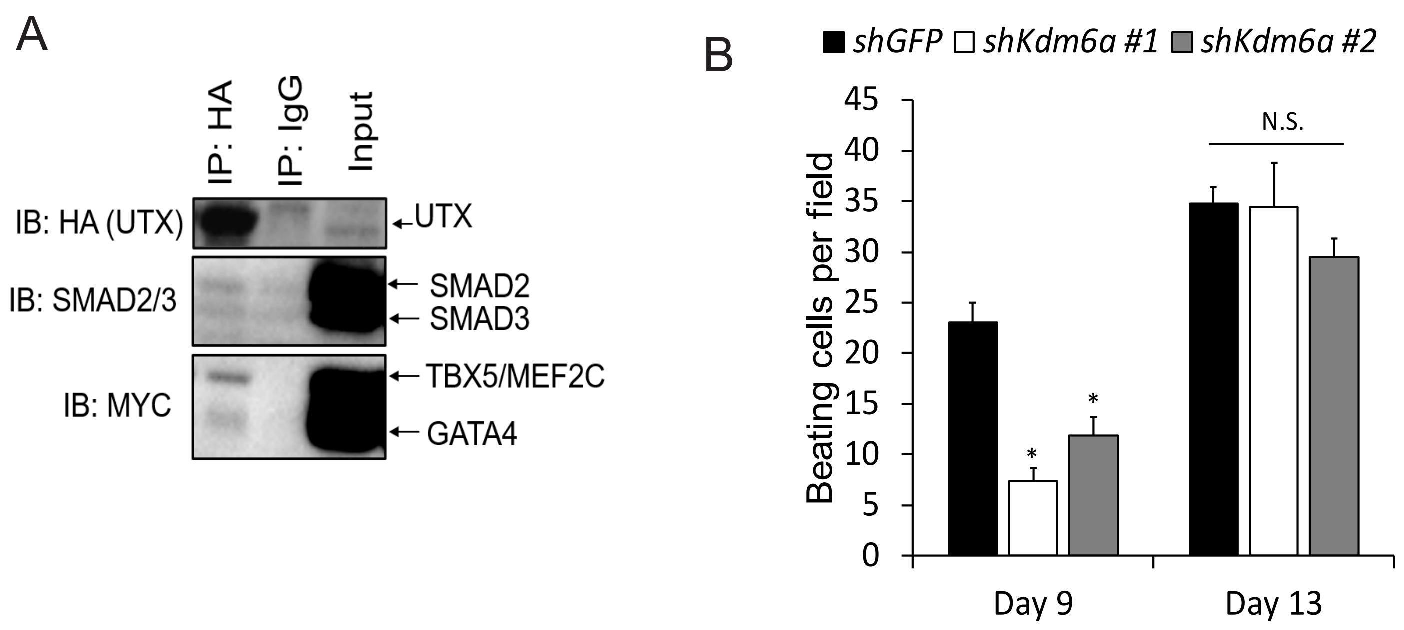
A
IP: HA
IP: IgG
Input
IB: HA (UTX)
UTX
IB: SMAD2/3
SMAD2
SMAD3
IB: MYC
TBX5/MEF2C
GATA4
B
shGFP
shKdm6a #1
shKdm6a #2
Beating cells per field
45
40
35
30
25
20
15
10
5
0
N.S.
*
*
Day 9
Day 13

Figure S3

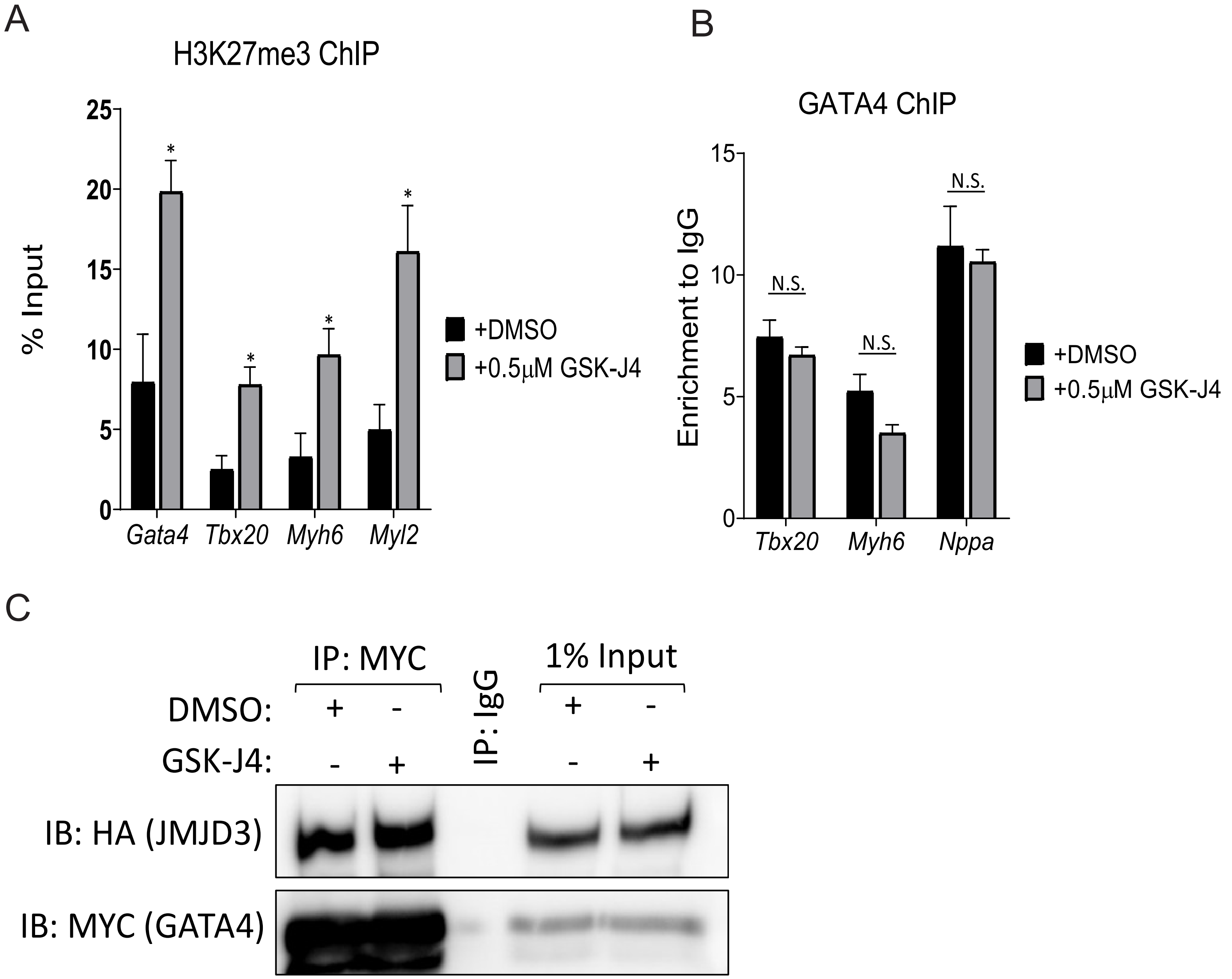

A
H3K27me3 ChIP
% Input
25
20
15
10
5
0
*
Gata4
Tbx20
Myh6
Myl2
+DMSO
+0.5μM GSK-J4
B
GATA4 ChIP
Enrichment to IgG
15
10
5
0
N.S.
Tbx20
Myh6
Nppa
+DMSO
+0.5μM GSK-J4
C
IP: MYC
IP: IgG
1% Input
DMSO: + - + -
GSK-J4: - + - +
IB: HA (JMJD3)
IB: MYC (GATA4)

### Figure S4

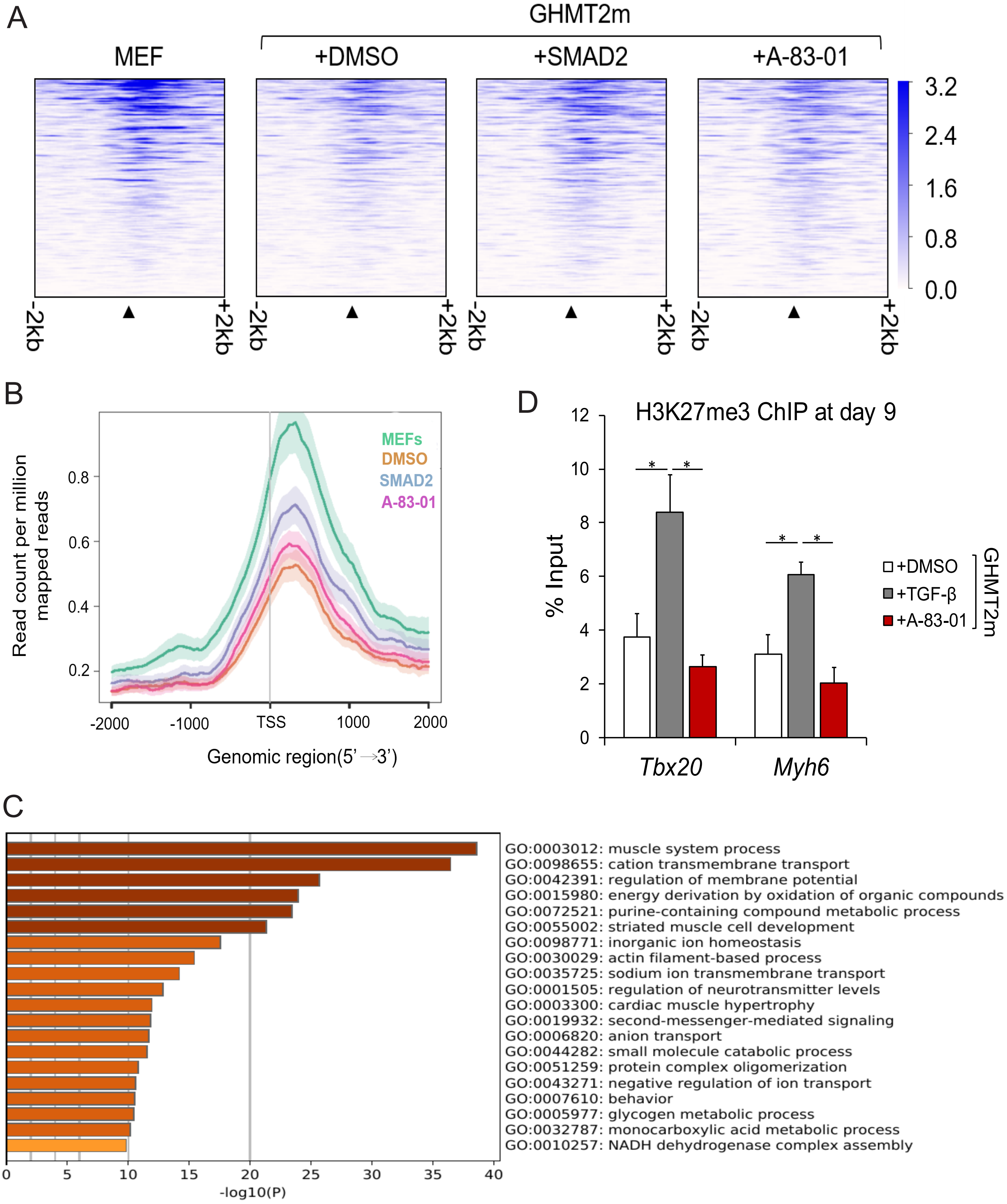
A
GHMT2m
MEF
+DMSO
+SMAD2
+A-83-01
3.2
2.4
1.6
0.8
0.0
-2kb
+2kb
B
Read count per million mapped reads
MEFs
DMSO
SMAD2
A-83-01
0.8
0.6
0.4
0.2
-2000
-1000
TSS
1000
2000
Genomic region(5'→3')
D
H3K27me3 ChIP at day 9
% Input
12
10
8
6
4
2
0
*
+DMSO
+TGF-β
+A-83-01
GHMT2m
Tbx20
Myh6
C
GO:0003012: muscle system process
GO:0098655: cation transmembrane transport
GO:0042391: regulation of membrane potential
GO:0015980: energy derivation by oxidation of organic compounds
GO:0072521: purine-containing compound metabolic process
GO:0055002: striated muscle cell development
GO:0098771: inorganic ion homeostasis
GO:0030029: actin filament-based process
GO:0035725: sodium ion transmembrane transport
GO:0001505: regulation of neurotransmitter levels
GO:0003300: cardiac muscle hypertrophy
GO:0019932: second-messenger-mediated signaling
GO:0006820: anion transport
GO:0044282: small molecule catabolic process
GO:0051259: protein complex oligomerization
GO:0043271: negative regulation of ion transport
GO:0007610: behavior
GO:0005977: glycogen metabolic process
GO:0032787: monocarboxylic acid metabolic process
GO:0010257: NADH dehydrogenase complex assembly
0
5
10
15
20
25
30
35
40
-log10(P)

Figure S5

A

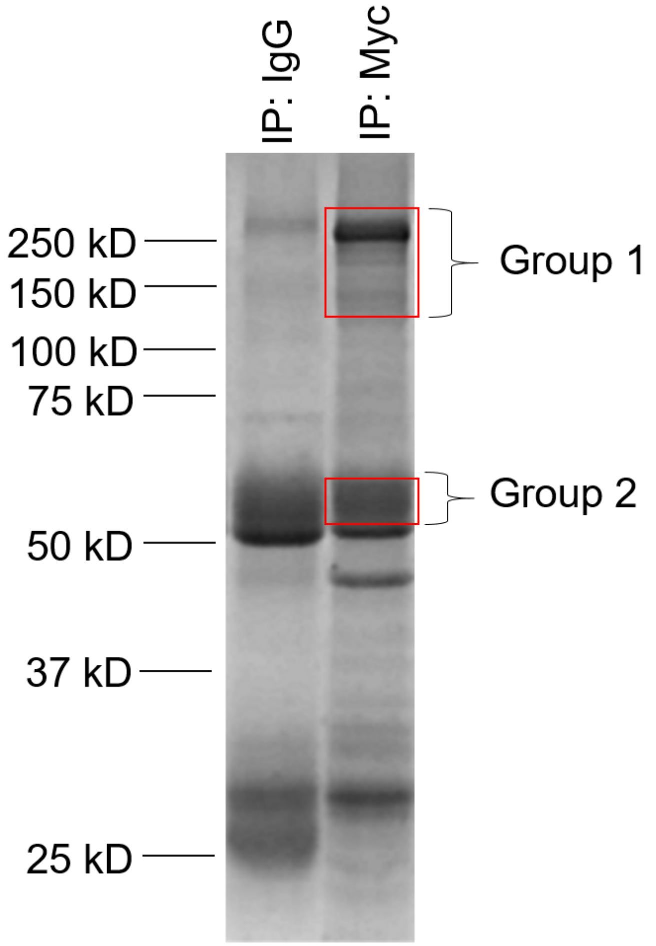


B

**Group 1**

| | |
|---|---|
| DHX9 | THOC2 |
| MBB1A | **BAF170** |
| PRP8 | HELZ2 |
| U520 | RFC1 |
| NUCL | RRP12 |
| RRP5 | PDS5A |
| DDX21 | NUP98 |
| RENT1 | SP16H |
| SNF2H | NOC3L |
| TRIPC | SRRM2 |
| BAZ1B | APC1 |
| TOP2B | ZFR |
| TOP1 | INT7 |
| HNRPU | SPB1 |
| INT1 | SLTM |
| NAT10 | CHD4 |
| ILF3 | **BAF155** |
| SF3B1 | BAZ1A |
| NOP2 | **BAF180** |
| U5S1 | CEBPZ |
| NU160 | MED23 |
| TIF1B | **BRM** |
| UTP20 | MED24 |
| NU133 | KDM1A |
| NU188 | NSD1 |
| MATR3 | CHD2 |
| HNRL2 | KMT2A |
| EDC4 | MED12 |
| TRRAP | KDM5B |
| PRP6 | INO80 |
| **BRG1** | MED14 |
| SNF2L | PHF2 |
| DDX54 | BAZ2A |
| CDC5L | DNMT1 |
| NU155 | **ARID1A** |

**Group 2**

| |
|---|
| GATA4 |
| TBX5 |
| MEF2C |

**Spectral Counts**

| |
|---|
| <10 |
| 10 to 25 |
| 25 to 50 |
| >50 |

Figure S6

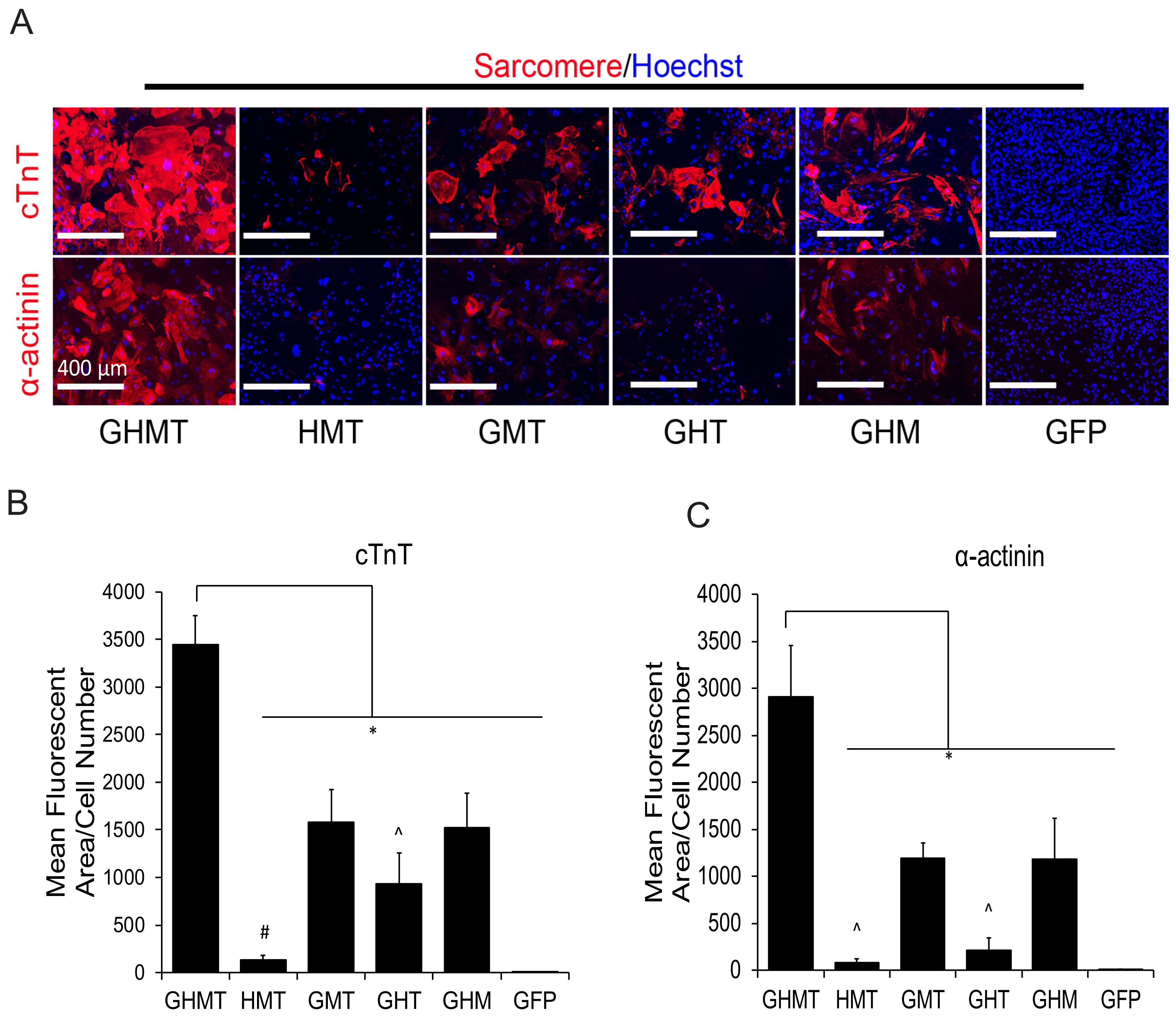

A
Sarcomere/Hoechst
cTnT
α-actinin
400 μm
GHMT
HMT
GMT
GHT
GHM
GFP
B
cTnT
Mean Fluorescent Area/Cell Number
4000
3500
3000
2500
2000
1500
1000
500
0
*
#
^
GHMT
HMT
GMT
GHT
GHM
GFP
C
α-actinin
Mean Fluorescent Area/Cell Number
4000
3500
3000
2500
2000
1500
1000
500
0
*
^
^
GHMT
HMT
GMT
GHT
GHM
GFP

Figure S7

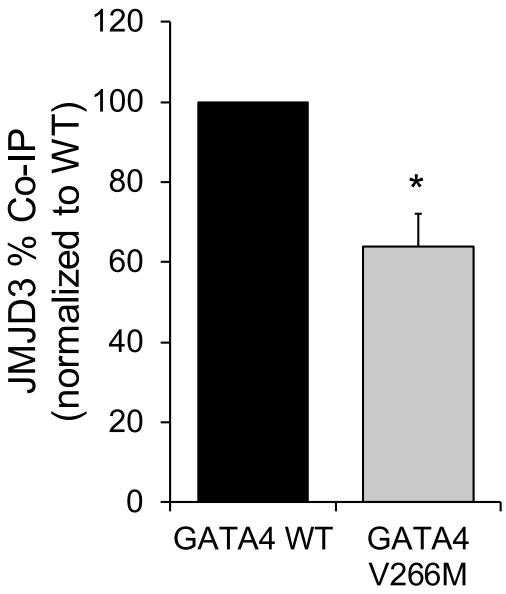

120
100
80
60
40
20
0
JMJD3 % Co-IP
(normalized to WT)
*
GATA4 WT
GATA4
V266M

Figure S8

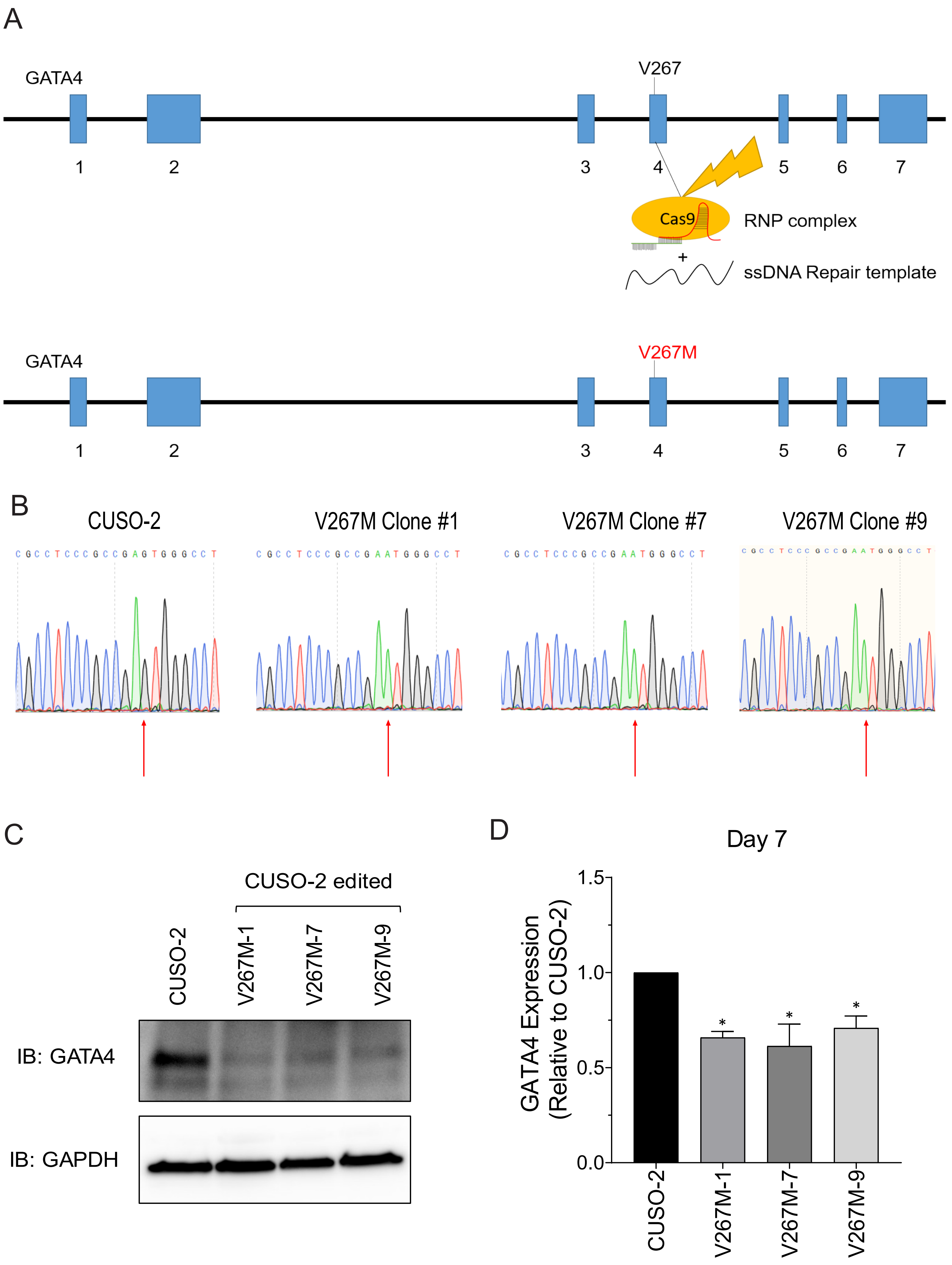
A
GATA4
V267
1
2
3
4
5
6
7
Cas9
RNP complex
+
ssDNA Repair template
GATA4
V267M
1
2
3
4
5
6
7
B
CUSO-2
V267M Clone #1
V267M Clone #7
V267M Clone #9
C G C C T C C C G C C G A G T G G G C C T
C G C C T C C C G C C G A A T G G G C C T
C G C C T C C C G C C G A A T G G G C C T
C G C C T C C C G C C G A A T G G G C C T
C
CUSO-2 edited
CUSO-2
V267M-1
V267M-7
V267M-9
IB: GATA4
IB: GAPDH
D
Day 7
GATA4 Expression
(Relative to CUSO-2)
1.5
1.0
0.5
0.0
*
*
*
CUSO-2
V267M-1
V267M-7
V267M-9

#### Supplemental Figure Legends

**Figure S1: Inhibition of canonical TGF-β signaling promotes cardiac reprogramming**

**A)** Beating cell counts per field (0.89 $mm^2$) in Day 10 GHMT2m-reprogrammed cells co-transduced with GFP or SMAD2. Reprogrammed cells were maintained in 0.5 µM A-83-01 starting at Day 3. N = 3 per group. **B)** Immunoblot of p-SMAD2 and Total SMAD2/3 from whole cell extracts of Day 12 GHMT2m-reprogrammed cells co-transduced with GFP or SMAD2. Reprogrammed cells were maintained in vehicle (DMSO) or 0.5 µM A-83-01 starting at Day 3.

**Figure 2S: UTX weakly associates with GHMT and SMAD2/3 and delays cardiac reprogramming kinetics.**

**A)** Co-IPs using nuclear lysates harvested from HEK293T cells transfected with MYC-GHMT and HA-UTX. Following Co-IP with anti-HA, proteins were resolved by SDS-PAGE and immunoblotted for MYC, HA, and SMAD2/3. **B)** Beating cell counts per field (0.89 $mm^2$) from Day 10 and 13 GHMT2m-reprogrammed cells co-infected with shGFP or two different shRNAs targeting *Kdm6a*. Reprogrammed cells were maintained in 0.5 µM A-83-01 starting at Day 3. N = 3 per group.

Data are presented as mean ± SEM. * $p<0.05$ by one-way ANOVA with Tukey's multiple comparison test vs the shGFP group.

**Figure S3: GSK-J4 prevents demethylation of H3K27me3 but does not disrupt recruitment of GATA4 to target genes or binding of GATA4 to JMJD3.**

**A)** H3K27me3 levels at cardiac gene promoters *Gata4, Tbx20, Myh6,* and *Myl2* from Day 7 GHMT2m-reprogrammed cells. Reprogrammed cells were treated with DMSO or 0.5 μM GSK-J4 between Days 3 and 5 and maintained in 0.5 μM A-83-01 starting at Day 3. Data are presented as percentage input. N=4 per group. **B)** GATA4 levels at cardiac gene promoters *Tbx20, Myh6,* and *Nppa* from Day 5 GHMT2m-reprogrammed cells. Reprogrammed cells were treated with DMSO or 0.5 μM GSK-J4 between Days 3 and 5 and maintained in 0.5 μM A-83-01 starting at Day 3. Data are presented as enrichment to IgG control. N=3 per group. **C)** Co-IPs using nuclear lysates harvested from HEK293T cells co-transfected with MYC-GATA4 and HA-JMJD3. Transfected cells were treated with DMSO or 0.5 μM GSK-J4 for 24 hours prior to lysis. Following Co-IP with anti-MYC, proteins were resolved by SDS-PAGE and immunoblotted for HA and MYC.

All data shown as mean ± SEM. * $p<0.05$ by Student's *t* test.

**Figure S4: Basal TGF-β signaling delays H3K27me3 demethylation kinetics.**

**A and B)** ChIP-seq density heatmap of H3K27me3 signal (A) and average peak enrichment plots (B) in Day 7 GHMT2m-reprogrammed cells with indicated TGF-β pathway manipulation and undifferentiated MEFs at ± 2kb from annotated transcription start sites (TSS) of genes listed in the Regulation of Heart Contraction Gene Ontology term (GO:0008016). **C)** Gene Ontology enrichment analysis of biological processes for >2-fold upregulated genes between GHMT2m-reprogrammed cells treated with A-83-01 and undifferentiated MEFs. Data were downloaded from the NCBI Gene Expression Omnibus, GSE71405. **D)** H3K27me3 levels at cardiac gene promoters *Tbx20* and *Myh6* from Day 9 GHMT2m-reprogrammed cells treated with DMSO, 5 ng/mL TGF-β1, or 0.5 μM A-83-01. Data are presented as percentage input. N=4 per group. Data are presented as mean ± SEM. * $p<0.05$ by one-way ANOVA with Tukey's multiple comparison test at specified comparisons.

**Figure S5: GATA4 is the most critical factor for GHMT-mediated reprogramming**

**A)** Co-IPs from nuclear lysates prepared from D5 GHMT2m-reprogrammed MEFs. Following Co-IP with anti-MYC or anti-IgG, lysates were resolved by SDS-PAGE. Acrylamide gel was stained with SPYRO Ruby to detect proteins. Individual bands (boxed regions) were excised and proteins were identified by mass spectrometry. **B)** Table of hits identified by mass spectrometry arranged by total spectral counts. Proteins listed in red indicate core components of the SWI/SNF complex.

**Figure S6: GATA4 is the most critical factor for GHMT-mediated reprogramming**

**A)** Representative fluorescent images of Day 12 MEFs reprogrammed with GHMT or GHMT-1 reprogramming factors. Cells transduced with GFP served as a negative control. Reprogrammed cells were immunostained with sarcomere proteins cardiac troponin T (cTnT) or α-actinin (Red) and co-stained with Hoechst for nuclei (Blue). Scale bar = 400 μM. **B and C)** Quantification of mean fluorescent area per field of cardiac markers cTnT (B) or α-actinin (C), normalized to the number of cells per field. 5 fields of view per dish were collected across 3 individual experiments per group to analyze expression of cardiac sarcomere proteins. Data are presented as mean ± SEM. *, ^, # $p<0.05$ by one-way ANOVA with Tukey's multiple comparison test vs all groups (*), GMT and GHM (^), or GMT, GHT, and GHM (#).

**Figure S7: GATA4 interacts with JMJD3 via a short linker between Zinc Finger domains including residue V266**

**A)** Quantification of interactions between JMJD3 and GATA4 shown in Figure 5E. N=3 per group. * $p$<0.05 by Student's $t$ test.

**Figure S8: GATA4 V267M disrupts iPS differentiation of cardiomyocytes**

**A)** CRISPR-Cas9 genome editing targeting strategy to generate GATA4 V267M iPSC lines. The crRNA and tracrRNA complex was incubated with Cas9 protein to form the RNA protein (RNP) complex. Single stranded DNA (ssDNA) carrying the GATA4 V267M point mutation was added to the parental WT GATA4 iPS line (CUSO-2) along with the RNP complex and electroporated. Cells were recovered and individual clones were selected and expanded. **B)** iPS clones were selected and homozygous GATA4 V267M positive clones were identified by Sanger sequencing. **C and D)** Western blot (C) and quantification (D) of GATA4 protein levels in Day 5 WT and V267M GATA4 iPS-CMs. Following whole cell lysis, proteins were resolved by SDS-PAGE and immunoblotted for GATA4 and GAPDH.

Data are presented as mean ± SEM. * $p$<0.05 by one-way ANOVA with Tukey's multiple comparison test vs the WT GATA4 (CUSO-2) line.

**Movies S1-S5**

Representative field of view of spontaneous beating of Day 13 iCMs transduced with GHMT2m co-expressing GFP and treated with DMSO (S1), 5 ng/µL TGF-β (S2), or 0.5 µM A-83-01 (S3), or co-expressing SMAD2 (S4) or SMAD7 (S5). Cells were visualized with a 10X microscope objective lens.
